# Recurrent phases of strict protein limitation inhibit tumor growth and restore lifespan in a *Drosophila* intestinal cancer model

**DOI:** 10.1101/2023.01.18.524563

**Authors:** Roxana M. Pfefferkorn, Benedikt M. Mortzfeld, Christine Fink, Jakob von Frieling, Judith Bossen, Daniela Esser, Christoph Kaleta, Philip Rosenstiel, Holger Heine, Thomas Roeder

## Abstract

Diets that restrict caloric or protein intake offer a variety of benefits, including decreasing the incidence of cancer. However, whether such diets pose a substantial therapeutic benefit as auxiliary cancer treatments remains unclear. We determined the effects of severe protein depletion on tumorigenesis in a *Drosophila melanogaster* intestinal tumor model, using a human RAF gain-of-function allele. Severe and continuous protein restriction significantly reduced tumor growth but resulted in premature death. Therefore, we developed a diet in which short periods of severe protein restriction alternated cyclically with periods of complete feeding. This nutritional regime reduced tumor mass, restored gut functionality, and normalized the lifespan of oncogene-expressing flies to the levels observed in healthy control flies. Furthermore, this diet reduced the chemotherapy-induced stem cell activity associated with tumor recurrence. Transcriptome analysis revealed long-lasting changes in the expression of key genes involved in multiple major developmental signaling pathways. Overall, the data suggest that recurrent severe protein depletion effectively mimics the health benefits of continuous protein restriction, without undesired nutritional shortcomings. This provides seminal insights into the mechanisms of the transcriptomic memory effect required to maintain the positive effects of protein restriction throughout the phases of a full diet.

## Introduction

Colorectal cancer (CRC) is the 3^rd^ most common cause of cancer-related deaths worldwide, accounting for approximately 1,8 million deaths and more than two million new cases annually (1, 2). CRC cases are expected to increase over the next decade because of the high incidence of major risk factors, such as obesity, physical inactivity, and tobacco and alcohol abuse (1, 3–5). Intestinal stem cells (ISC), which tend to accumulate mutations because of their proliferative properties and longevity, are particularly important for the development of CRC. ISCs can develop into tumor stem cells when their signaling pathways involved in proliferation control, such as the EGFR signaling pathway, are affected. Mutations in oncogenes, the products of which are active downstream of EGFR, are among the most common driver mutations in nonfamilial CRC. Mutations in KRAS, BRAF, PTEN, and PIK3 are particularly prominent (6, 7).

Signaling pathways that play a role in cell proliferation are frequently involved in the regulation of cell metabolism. This dual function indicates that cell proliferation is closely related to the availability of energy and metabolites. Consequently, changes in diet that affect energy and/or metabolite supply can have a direct impact on proliferation as well as on tumor development and growth. This link is supported by epidemiological studies showing that obesity, diabetes, and dietary habits are correlated with tumor development (3, 8). Based on this interconnection, particular dietary interventions have been used in cancer therapy, although the data supporting such an approach are relatively sparse. For instance, it has been suggested that the metabolism of cancer cells can be altered by the withdrawal of carbohydrates while granting caloric intake through fats, which can limit the proliferative potential of tumors (9–11) through the production of ketone bodies (11). This effect can be obtained by restricting the total energy supply (resulting in an overall reduction in caloric intake) or by restricting certain metabolites (particularly proteins) to specifically interfere with the signaling pathways in which relevant driver mutations are active. Reductions in overall calorie intake have shown largely consistent inhibitory effects on cancer incidence and progression in various animal models (12–14). Protein limitation has been shown to reduce the growth of multiple tumors *in vitro* and in xenograft models, and to reduce the incidence of multiple types of cancer (15–17), pointing to a potential therapeutic strategy.

The degree of caloric or protein restriction is crucial for balancing the needs of normal cellular metabolism with the desired cellular reprogramming that results in tumor inhibition. Although complete calorie intake restriction by fasting would have the greatest effect on tumor cells, this is not a feasible strategy. Instead, intermittent fasting regimes have been developed in which phases of severe calorie reduction alternate with normal nutrition. The restrictive phase can be achieved through complete starvation, but patient compliance may be more feasible with a fasting mimicking diet (FMD). Mouse models of diabetes and multiple sclerosis have shown considerable physiological improvements upon intermittent FMD, including extended longevity, increased cognitive performance, improved immune function, and reduced overall morbidity (18–21). In humans, a clinical trial with recurrent FMD resulted in reduced IGF-1 and blood glucose levels and lowered blood pressure (22). The first clinical trial to determine the effect of FMD in cancer patients showed promising preliminary results (23).

Complex diet plans that restrict total caloric intake are now increasingly being studied experimentally, but the equally promising reduction in protein intake is addressed less frequently. This is mainly because although protein restriction inhibits tumor growth, adequate protein intake following the ESPEN guidelines for tumor patient diets (24) makes it difficult to implement this strategy as a means of tumor therapy. To overcome this apparent dilemma, strategies have been adopted in which individual essential and nonessential amino acids are selectively depleted (25, 26). To use the primary concept of reduced total protein intake, despite the difficulties described above, dietary regimes in which phases of strict protein restriction alternate with phases of a normal diet are suitable. Using a *Drosophila melanogaster* model, we have already shown that such a strategy exhibits a robust general life-prolonging effect in the fruit fly (27) and suggest its utility as an auxiliary cancer therapy.

*Drosophila* models have been shown to be highly applicable for investigating essential aspects of human tumor development (28, 29), and concrete therapeutic suggestions can be made under specific conditions (30). We used the *Drosophila* CRC model established by Markstein (31) to investigate the effects of a reduced protein diet on tumor growth and host survival. The model allows for tight spatiotemporal determination of RAF gain-of-function (RAF^gof^) oncogene expression, whereas co-expression of luciferase and green fluorescent protein (GFP) allows for tumor traceability and quantification. First, we determined the most effective degree of protein reduction for tumor growth inhibition, followed by mechanistic investigations to reveal how these dietary interventions affected the interplay between tumor inhibition and life expectancy. The results presented here show that tumor size was significantly reduced in flies subjected to a strict protein depletion diet (PDD), whereas continuous moderate protein reduction was less effective. However, PDD results in an extensive loss of body mass and drastically reduces the lifespan of the flies. In contrast, a feeding regime in which phases of strict protein depletion are alternated with normal feeding reduces tumor size as effectively as PDD but restores life expectancy to normal levels. We identified a unique transcriptomic signature in flies subjected to recurrent protein depletion that provides crucial mechanistic insights into how such a diet provides long-term benefits through transcriptomic memory effects that mimic the effects of lifelong nutritional restrictions.

## Material and Methods

### Fly Husbandry

Homozygous flies were used with *esg-Gal4, UAS-GFP, tubulin-Gal80^ts^* on chromosome *2*, and *UAS-luciferase* on chromosome *3*, referred to as EGT;Luc2. These were crossed with homozygous *UAS-RAF^gof^* flies (RRID:BDSC_2033) to obtain the F1 generation with a temperature-inducible (29°C) tumor phenotype (EGT;Luc2 > RAF^gof^). Both parent fly lines were gifts from Markstein (31). As a control, EGT and Luc2 flies were crossed with *w^1118^* (RRID:DGGR_108479) wild-type flies (EGT;Luc2 > w^1118^). All stock fly lines and crossings were raised in cornmeal-based standard medium at 20°C. The F1 generation was shifted to 29°C 5-7 days post eclosion to induce the expression of *UAS*-dependent genes. Only the mated female flies were used in the assays. Flies from multiple independent breedings were mixed and randomly assigned to treatment groups. Experiment were unblinded. No prior power analysis was performed, our sample sizes are in line with or exceed previous studies in this field.

All animal experiments were performed in compliance with the regulations and ordinances for ethics and animal protection by the local authorities of the land Schleswig-Holstein (MELUND, Kiel, Germany).

### Fly Food and feeding regimes

All assays were conducted using low-melt fly food (31) containing 7% corn syrup and 1.5% agar, supplemented with 0.1% propionic acid and 0.3% nipagin to prevent microbial growth. The amount of protein varied in Bacto yeast extract (YE). The nutritious control diet (CD) contained 2% YE. After comparing the contents of 0.1, 2, 5, and 7.5% YE, a protein depletion diet (PDD) containing 0.1% YE was chosen.

Three feeding regimes with recurrent protein restrictions were tested. Regime 1 involved an initial phase of five days on PDD, followed by repeated cycles of three days on CD and four days on PDD. The same cycling regime was used as Regime 2; however, the initial phase on PDD only lasted for 3 d. Regime 3 started with 3 d on CD, followed by cycles of 3 d on PDD and 4 d on CD.

### Vibratome Sectioning and microscopy

The flies were dissected on specified days, typically 3–10 d after induction at 29°C. After removal of the head and anal plate, the body was fixed in 4% paraformaldehyde for 12 h. Following embedding in 7% agarose, sections of 100 μm were produced using a vibratome (80 Hz, 0.5 mm amplitude). When indicated, the first sections of the thorax containing the proventriculus and the R1 midgut region were discarded to ensure that only the anterior R2 midgut was assessed. Actin was stained with phalloidin (1:1000 in PBS, 10 min incubation) and the nuclei were stained with DAPI following standard procedures. EGT^+^cells were labeled with GFP. Gut and lumen volumes were measured in cross sections using ImageJ 1.49 v (RRID:SCR_003070).

### Luciferase Assay

Luciferase activity was determined using the One Glo Luciferase Assay System (Promega, Walldorf, Germany, #E6110). For each replicate, three flies were collected in 150 μL Glo Lysis Buffer and homogenized using a bead mill homogenizer (Biolab Products, Bebensee, Germany) for 2 min at 3.25 m/s. The suspension was centrifuged for 3 min at 12,000 × g, and 50 μL of the supernatant was mixed with an equal volume of the substrate. Luciferase signals were detected using a Tecan plate reader (Tecan, Crailsheim, Germany).

### Gut Integrity

Age-matched fly populations (10 individuals per replicate) were incubated at 29°C while feeding on CD supplemented with Brilliant Blue FCF food dye (Carl Roth, Karlsruhe, Germany, #2981.2) for three days. Leaky gut is indicated by a blue dye distributed throughout the body (32).

### Metabolic Rate Assay

Basic metabolic rate was determined as previously described (33). The chamber was set up by using multiple fly respirometers. Each respirometer was composed of microcapillary air tightly attached to a 1 ml micropipette tip. Slightly moist soda lime was placed between the two cotton plugs inside the tip. Per replicate three flies were placed in each respirometer, which was then sealed with plasticine and placed in a TLC chamber filled with water containing the Brilliant Blue FCF food dye for visual inspection. Two respirometers without flies were used as the atmospheric controls. As the CO_2_ produced by the flies was absorbed by the soda lime, the subsequent low-pressure was quantified by measuring the ascent of water inside the microcapillaries over two hours. The CO_2_ production was calculated as μL/h per fly.

### Fly survival

Age-matched fly populations (30 flies per replicate) were incubated at 29°C with the indicated food regimes and dead and live flies were counted every 2^nd^ day. The food was changed at least every 3^rd^ day. The populations were monitored until all flies died.

### Fecal Output Measurements

Aslope fly vials were filled with a medium supplemented with Brilliant Blue FCF food dye, and three flies per group and replicate were trapped with the food using a microscope coverslip and vial plug. Following incubation at 29°C, cover slips with defecation spots were collected, scanned, and analyzed using T.U.R.D. software (34) with the settings offset = 15, min size = 50, and max size = 1000. Based on this, total fecal output, number of fecal spots, and spot size were calculated.

### Feeding Behavior

The food interaction rate of the flies was measured using the FLIC system (35). It was equipped with 12 individual food arenas per unit, where each arena contained a food reservoir containing 5% glucose solution surrounded by a signal pad. Per replicate single flies from each treatment group were placed in the arena and monitored for 2 h. The fly must touch the signal pad to reach the food source, thus closing the electrical circuit. By recording the electric current, the exact number of food interactions can be recorded for each fly.

### Body Composition

Fly weight was assessed for three flies per replicate following collection in pre-weighed 2 ml screw cap tubes using a microbalance (Ascuro, Lörrach, Germany), addition of 1 ml of 0.1% Triton-X 100 in PBS, and homogenization. The homogenate was centrifuged for 3 min at 12,000 × g and the supernatant was used for body fat and protein analyses. Body fat content was measured as previously described (36). The homogenate was heated to 70°C for 5 min, and 50 μL was transferred to a 96-well plate, to which 200 μL of triglyceride reagent (Thermo Fisher Scientific, Germany, #T2449) was added per well. The absorbance at 540 nm (A_540_) was measured after 30 min of incubation at 37°C using a plate reader (Labexim Products, Lengau, Austria). The protein content was quantified using the Pierce™ BCA Protein Assay Kit (Thermo Fisher Scientific, Dreieich, Germany, #23227) with 25 μL of supernatant mixed with 200 μL of reagent, and the A_540_ was determined after 30 min of incubation at 37°C.

### Dechorionation and recolonization

To raise germ-free flies, eggs were collected from a 10% apple juice agar plate glazed with a yeast/water solution and dechorionated with 6% NaClO for 5 min. The dechorionated eggs were washed with 70% ethanol for 2 min and then three times with sterile water for 5 min. Disinfected eggs were hatched in sterile vials containing standard cornmeal-based fly food. Recolonization with a mixture of *Lactobacillus plantarum^WJL^, Lactobacillus brevis^EW^, Acetobacter pomorum, Commensalibacter intestini^A911T^*, and *Enterococcus faecalis* (all received as gifts from Carlos Ribeiro, Champalimaud Center for the Unknown, Lisbon, Portugal) was performed as previously described (37).

### Characterization of the intestinal bacterial microbiome

For microbiome analysis, oncogene-bearing and control flies were co-cultivated in separate culture flasks with gauze to provide an interface for bacterial exchange. The flies were then separated for induction at 29°C with their respective food sources for 10 d. The midguts of five individuals per replicate were dissected, transferred to sterile S2 medium, homogenized, and DNA extracted using a DNeasy Blood & Tissue Kit (Qiagen, Hilden, Germany, #69504). DNA was eluted in 100 μL of sterile AE buffer and stored at −20°C until sequencing. The bacterial variable regions 1 and 2 of 16S rRNA were amplified as previously described (38). The amplicons were sequenced using an Illumina MiSeq with 2 × 300 bp paired-end sequencing. After assembling the reads using SeqPrep, chimers identified using ChimeraSlayer (39) and manually checked, were removed. The sequences were further analyzed using QIIME 1.9.0 (40). The specific association of bacterial taxa with one of the conditions was tested using linear discriminant analysis (LDA) Effect Size, as previously described (41). To measure beta diversity and compare the groups, Principal Coordinate Analysis (PCoA) was performed using the beta_diverity.py script in Qiime.

### Transcriptome analysis

For RNA isolation, 15 midguts (taken from flies 13 days after tumor induction) were dissected per replicate and incubated in TRIzol for 5 min at room temperature. RNA was isolated using chloroform/ethanol extraction. RNA-seq libraries were prepared using the TruSeq Stranded Total RNA Library Prep Kit (Illumina, SanDiego, USA). After cluster generation on cBot 2 (Illumina, SanDiego, USA) using the HiSeq 3000/4000 SR Cluster Kit (Illumina, SanDiego, USA), sequencing was performed using the HiSeq 3000/4000 SBS Kit (50 cycles, Illumina, SanDiego, USA) on an Illumina HiSeq 3000/4000 (Illumina, SanDiego, USA). After quality filtering, the reads were mapped against the *Drosophila melanogaster* reference BDGP6 using the TopHat v2.1. (42) (RRID:SCR_013035) with a sensitive option. The read counts per transcript were calculated with the Python (RRID:SCR_001658) script HTSeq v0.6.1p1 (43) (RRID:SCR_005514) using the mode ‘intersection-strict’ mode. This step was based on annotation BDFP6.92. P-values for differentially expressed genes were calculated using DeSeq2 v1.14 (44) (RRID:SCR_015687). Outliers were detected and removed per gene using the function ‘replaceOutliersWithTrimmedMean’. A heatmap was created in R using the function ‘heatmap.2’ (45). Hierarchical clustering was performed with the R function ‘hclust’ using Euclidean distances based on log-transformed expression data (46).

GO analysis was performed using the ShinyGO program package (RRID:SCR_019213) with an FDR cutoff of 0.05 and a selection of the top 20 pathways (47).

### Drug exposure

Adult flies were subjected to the indicated substances for three consecutive days. Afatinib, sorafenib, LiCl_2_, SP600125, DBZ, baricitinib, filgotinib, and oclacitinib were dissolved in 0.1% DMSO and added to CD fly food at a final concentration of 100 μM. Luciferase activity was measured as previously described.

### PH3 quantification

Adult *w^1118^* flies were subjected to 100 μM paclitaxel (dissolved in 0.1% DMSO and added to the feed) for three consecutive days. The intestines were dissected and fixed in 4% paraformaldehyde for 1 h before blocking with 5% NGS and staining with rabbit-PH3 primary antibody (Cell Signaling Technology, Leiden, Netherlands, #3377) overnight, followed by detection using anti-rabbit AF488 as a secondary antibody. Images were obtained using an Axio Imager.Z1 or LSM 880 (Zeiss, Jena, Germany) and PH3^+^ cells were counted manually.

### Statistical Analysis

Statistical methods to predetermine sample sizes were not used but the sample sizes are similar to or higher than those used in previous studies (36, 48). Specific approaches to randomly allocate samples to groups were not used and the experiments were not performed in a blinded design. No data were excluded from the analysis. Data were tested for normality using the Shapiro-Wilk test and for homogeneity of variance using the Levene test. Depending on the outcome, a pairwise t-test or Welch’s t-test was used for parametric data or the Wilcoxon ranksum test was performed for non-parametric data and datasets with low sample sizes. ANOVA was used when applicable. We used the false discovery rate to correct multiple tests. Data are either represented as boxplots or as the mean ± standard error of the mean (SEM, represented by error bars in the line graphs). Statistical significance was set at P < 0.05.

## Results

### Expression of RAF^gof^ in intestinal progenitor cells induces tumor growth and reduces lifespan

To induce stem cell tumors in the intestines of adult flies, a human RAF gain-of-function allele (UAS-RAF^gof^) was exclusively expressed in ISCs and enteroblasts using a temperature-sensitive esg^+^driver. The co-expression of luciferase and GFP in an identical spatiotemporal pattern in this model allows the assessment of tumor mass directly through microscopic evaluation and indirectly by measuring luciferase activity (31). As expected, the expression of RAF^gof^ in intestinal progenitor cells resulted in an increase in GFP-marked progenitor cells in the midgut compared to the control (Fig. 1A, upper panel). Thoracic cross sections revealed dense cell accumulation protruding into the intestinal lumen (Fig. 1A, lower panel). Morphological restructuring was dispersed throughout the entire intestine, with increased progenitor cell abundance in each investigated midgut region. Over a period of 10 days, the development of ISCs into enteroblasts was clearly visible (Fig. S1). The intestinal structure and phenotypic variability of the control and oncogene-expressing flies are shown in Fig. S2. Luciferase activity in these flies depends on progenitor cell abundance as well as on the strength of promoter activation. Induction of over-proliferation through the expression of RAF^gof^ in flies fed a fully nutritious control diet (CD) resulted in a four-fold increase in progenitor cell signal within three days of tumor induction (Fig. 1B). We assessed whether oncogene expression disturbed the intestinal barrier; however, this was not the case, as the gut integrity was not significantly impaired (Fig. 1C). As the control flies aged, their basic metabolism decreased, as measured by respiratory CO_2_ generation at rest. In contrast, the oncogene-expressing flies showed a more than two-fold higher resting metabolic rate, in line with the increased energy demand of highly proliferative tissue (Fig. 1D).

**Fig. 1:**
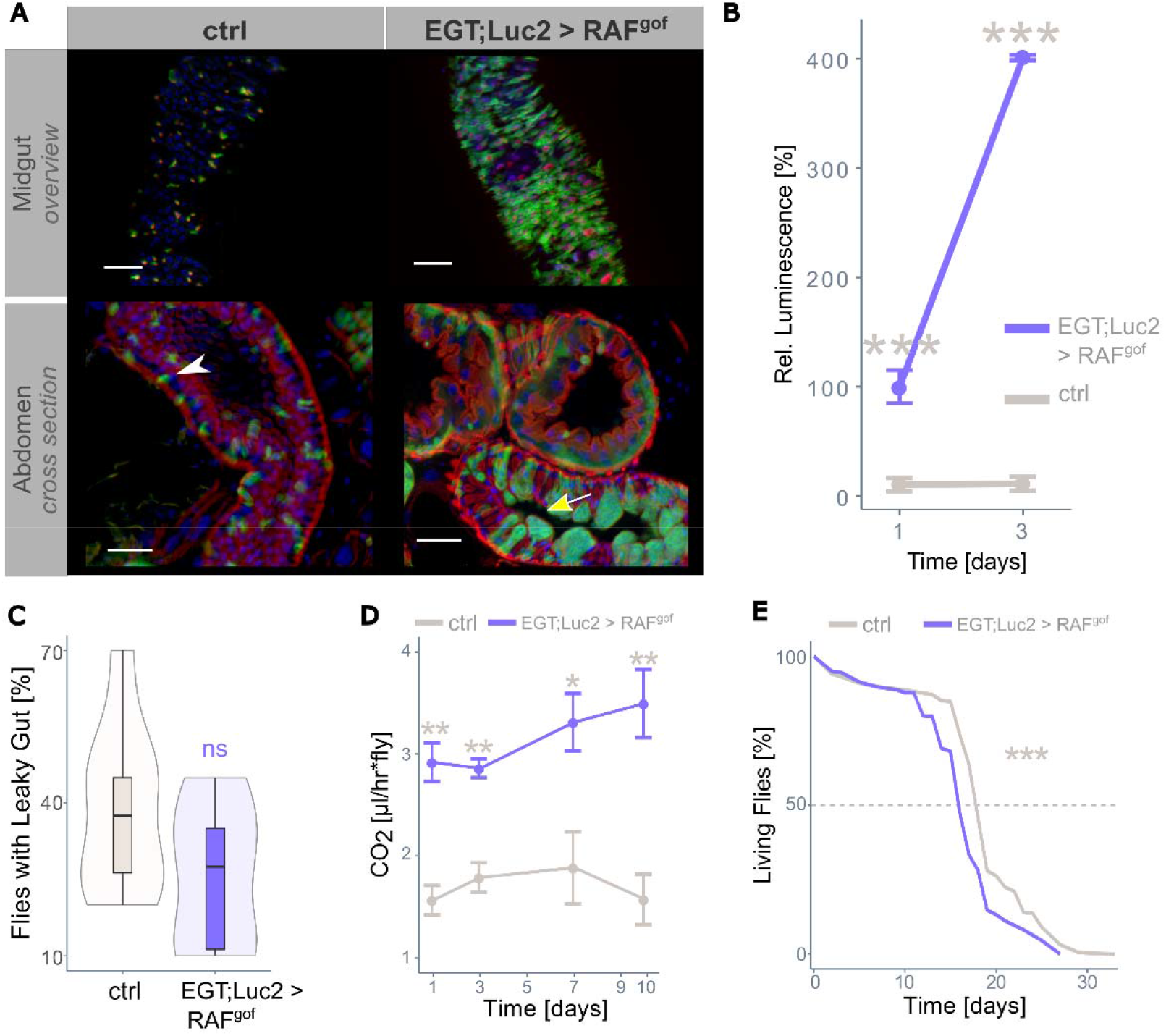
RAF^gof^ expression in intestinal stem cells and progenitor cells leading to tumor formation. (A): Overview of the posterior midgut (top panels) and cross section of the abdomen (lower panels) of control (ctrl, EGT;Luc2 > *w^1118^*) and tumor-bearing flies (EGT;Luc2 > RAF^gof^) after 3 days of oncogene induction (29°C). The white arrowhead indicates normal sized nuclei in controls and the yellow arrow enlarged nuclei in RAF^gof^ expressing flies. Note the cell clusters protruding into the lumen. Actin was stained with phalloidin (red) with additional staining of nuclei with DAPI (blue). Scale bar: 50 μm. (B) RAF^gof^ co-expressed luciferase signal in control flies and tumor-bearing flies (mean ± SEM, n=5-6). The level of luminescence measured after 24h of oncogene induction in the tumor-bearing flies is taken as 100%. (C) The gut integrity of healthy controls and tumor-bearing flies was not significantly affected after 3 days of oncogene induction (n=10). (D) Resting metabolic rate of control and tumor-bearing flies over time (mean ± SEM, n=5-10). (E) Lifespan of control and tumor-bearing flies at 29°C (n=450 flies per treatment). In all experiments flies were fed a normal control diet. Here and in all figures, significance is indicated as * p < 0.05; ** p < 0.01; *** p < 0.001; ns: not significant, with colors representing the partner for comparison

The lifespan of tumor-bearing individuals, independent of the host organism, tumor type, or location, is typically reduced proportionally to the tumor load and aggressiveness because a high tumor load can result in organ failure, whereas high tumor aggressiveness increases the risk of metastasis to distant organs. In our model, healthy control flies fed CD had a median lifespan of 19 days when reared at a permissive temperature (29°C), whereas ectopic expression of RAF^gof^ shortened the median lifespan to 16 days (Fig. 1E). Thus, overexpression of this oncogene significantly reduced the lifespan of the flies.

### Severe protein depletion reduces tumor load and reinstates gut functionality

The yeast extract (YE) content was varied in the diet of the flies to establish a protein-depleted diet (PDD) that could be used to assess its effect on tumor mass. Feeding the oncogene-bearing flies a diet containing various amounts of YE for three days revealed a strong correlation between dietary protein concentration and tumor growth, as quantified by luminescence (Fig. 2A). A severely protein-restricted diet containing only 0.1% YE resulted in a drastic reduction in tumor mass, which was used as the PDD in all subsequent experiments. After feeding the flies with PDD for three days, structural changes were visible in the dissected midguts and crosssections of their intestines (Fig. 2B; additional images are shown in Fig. S3). The dissected midguts showed a high abundance of GFP^+^ progenitor cells in oncogene-expressing flies fed on CD, whereas PDD drastically reduced this abundance, although the accumulation of progenitor cells and enlarged nuclei was still apparent (Fig. 2B, upper panel). Healthy control flies had very few esg^+^ cells in thoracic cross-sections, and oncogene-expressing flies fed on CD displayed an excessive intestinal mass with a solid layer of esg^+^ progenitor cells that pushed differentiated esg^-^ cells towards the gut lumen (Fig. 2B, lower panel, asterisk). In contrast, PDD feeding resulted in slightly more esg^+^ cells than healthy controls but lacked this superfluous cell layer (Fig. 2B). Sporadically, the PDD-treated flies displayed single esg^+^ cells detached from the basal membrane, which was not observed in control flies (results not shown).

**Fig. 2:**
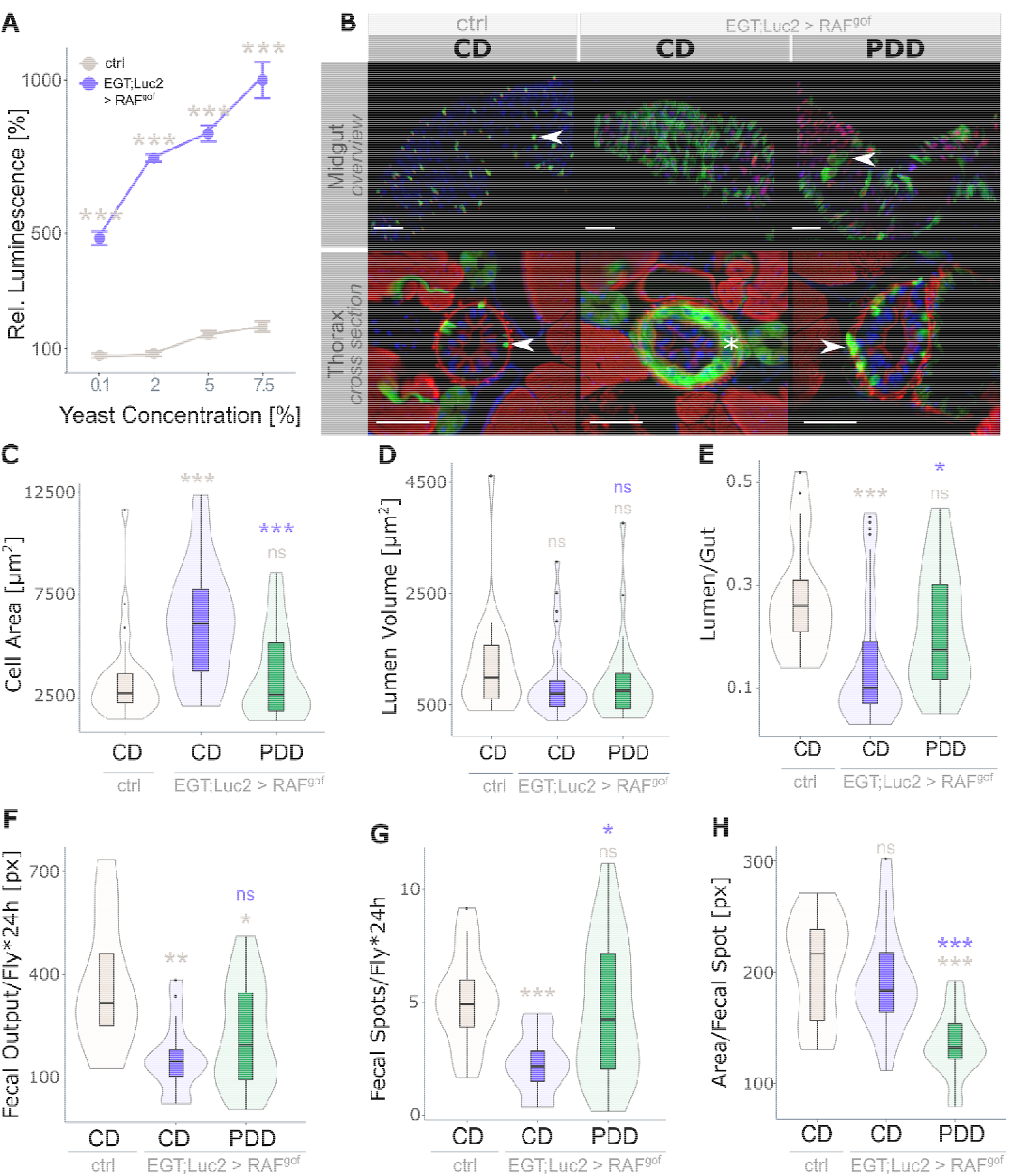
Strict protein depletion reduces tumor phenotype. (A) Effect of dietary protein in the form of yeast extract (YE) on midgut intestinal progenitor cell abundance in healthy controls and oncogene-expressing flies (n=8). Progenitor cell activity in control flies fed on CD (2% YE) is taken as 100%. (B) Overview of the midgut (upper panel) and thorax cross section (anterior midgut, R2, lower panel) of control flies fed on CD and of flies expressing RAF^gof^ fed on CD and on PDD containing 0.1% YE. Arrow heads point to GFP-marked esg^+^ cells attached to the basal membrane. The asterisk indicates the additional layer of esg^+^ cells in oncogene-expressing flies subjected to CD (scale bar: 50 μm). (C-E) Physiological measurements of the anterior R2 midgut (n=21-33) with (C) cell area per cross section, (D) lumen volume per cross section, and (E) lumen to gut ratio. (F) Quantification of the total fecal output in 24 h (n=9-10), with (G) fecal spot quantity and (H) size of individual fecal spots. All data were obtained after 3 days of oncogene induction at 29°C.

The effect of RAF^gof^-induced proliferation in the R2 region of the intestine was assessed using various physiological parameters. The cell area per cross-section was significantly higher in oncogene-expressing flies fed CD than in controls, but this was restored to normal levels on PDD (Fig. 2C). The volume of the gut lumen per cross-section was not significantly affected (Fig. 2D); however, the lumen-to-gut ratio indicated luminal obstruction in CD-fed flies expressing RAF^gof^, whereas the phenotype was rescued by PDD (Fig. 2E).

Because continuous PDD rescued the morphological changes induced by oncogene expression, we investigated the impact of PDD on gut functionality. The nutritional value of ingested food is highly dependent on the physiological state of the fly and epithelial ability to absorb and metabolize nutrients. To quantitatively assess this, the fecal output of healthy and RAF^ŝθf^-expressing flies was determined over a period of 24 h, which revealed that tumor induction led to a reduction in the total fecal output independent of diet (Fig. 2F). The total fecal output is the product of fecal spot abundance and spot size. The reduced output of tumor-bearing CD-fed flies could be attributed to the decreased number (Fig. 2G) of otherwise normally sized fecal spots (Fig. 2H). This indicates an impediment to digestive function as food passes through the gut, most likely caused by an excessive cellular mass protruding into the gut lumen. In contrast, the reduced fecal output of flies expressing PDD was the result of a high number of small individual fecal spots, indicating the regular digestion of less nutritious dense food.

### Prolonged severe protein depletion shortens lifespan

Continuous moderate protein restriction has been shown to have many benefits in healthy flies and can extend their lifespan by up to 50% (49). However, the severe protein restriction of PDD applied here led to a much shorter median lifespan of only nine days in RAF^gof^ expressing flies (Fig. 3A, green line) compared to 17 days when the flies were fed on CD and 19 days in control flies on CD (Fig. 3A, gray dashed line). The shortened lifespan was not attributable to tumor load, since the oncogene-expressing flies fed on PDD had morphologically mostly normal functioning intestines, and PDD also shortened the lifespan of the control flies by 7 days (Fig. S4). Food interactions were more frequent in RAF^gof^-expressing flies on the PDD than in CD-fed controls (Fig. 3B). The weight of the oncogene-expressing flies fed on CD showed minor fluctuations over time, which did not significantly differ from that of the controls on days 7 and 10, but their weight steadily declined on PDD (Fig. 3C). This severe loss of body mass is reflected in the loss of body fat and protein. The triacylglyceride levels of oncogene-expressing flies were not reduced when fed CD, but drastically decreased over time in flies subjected to PDD until approximately 25% body fat remained (Fig. 3D). Similarly, the protein content of tumor-bearing flies fed CD gradually normalized to that of control flies over time, whereas it severely decreased in PDD-fed flies (Fig. 3E). In conclusion, tumor-bearing flies fed a fully nutritious medium displayed mild alterations in body composition, whereas a prolonged, severely protein-restricted diet resulted in major depletion of body fat and protein and shortened the lifespan of the flies.

**Fig. 3:**
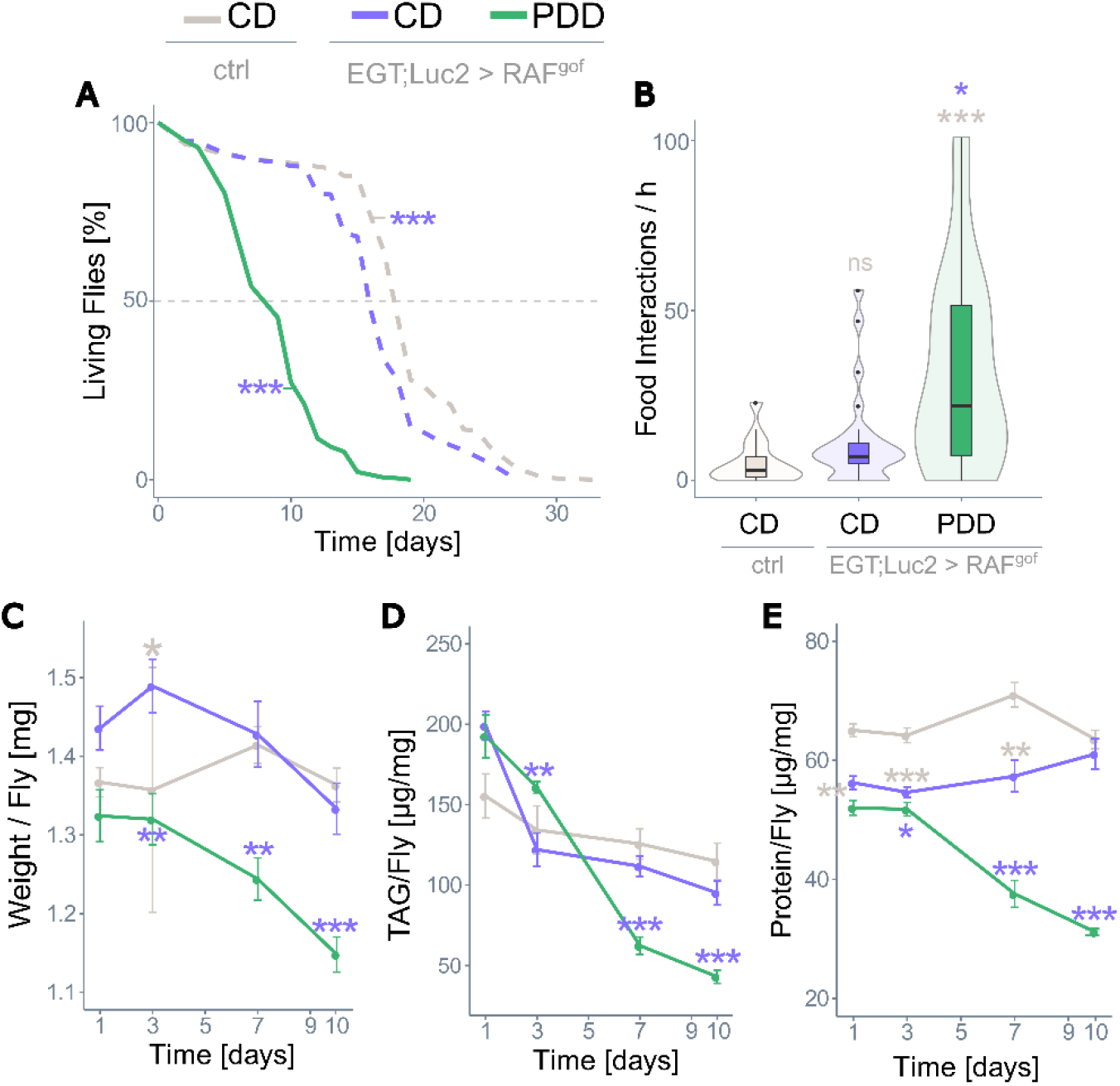
Prolonged protein depletion does not sustain life. (A) Lifespan of control flies on CD and of RAF^gof^ expressing flies on CD or PDD at 29°C (n=450 flies per treatment). (B) Food intake of control and RAF^gof^ expressing flies (n=22-29). (C) Body weight over time (n=14-17), (D) body fat as triacylglyceride (TAG) content (n=14-18), and (E) protein content (n=9-10) of control flies on CD and of RAF^gof^ expressing flies on CD or PDD at 29°C.

### Short cycles of severe protein depletion rescue the lifespan of oncogene-expressing flies

Although severe protein restriction effectively halted tumor growth, its negative effects on body composition and lifespan would prohibit its use as a therapeutic dietary intervention. Recent studies have suggested that diets with alternating phases of fully nutritious and moderately protein-restricted food can prolong *the Drosophila* lifespan (50) to an extent similar to lifelong dietary restriction (27). Therefore, we investigated the effects of such dietary regimes on longevity using three different feeding regimes that differed either in the duration of the initial feeding on PDD or the duration of the cycling phases. The first investigated diet (Regime 1) started with five days on PDD, followed by cycles of three days on CD and four days on PDD. The viability of flies during this feeding regime is shown in Fig. 4A, in which continuous diets of CD and PDD plus control flies on CD were added for comparison. While Regime 1 treatment rescued the overall lifespan of RAF^gof^ expressing flies to the control level, the initial phase of five days on PDD followed by three days on CD resulted in the death of 35-40% of flies (Fig. 4A). Despite this, the fly population had a median lifespan of 21 days, which was two days longer than that of the control (Fig. 4A). To minimize the lethal effect of the first eight days, in Regime 2, the initial PDD period was shortened to three days, while keeping the subsequent cycles the same as in Regime 1. This treatment rescued the median lifespan of the RAF^gof^-expressing flies without increasing their lethality during the first week of treatment (Fig. 4B). A third regime that started with three days of CD, followed by cycles of three days of PDD and four days of CD, also rescued the median lifespan (Fig. 4C). The maximum survival, calculated for the 10% longest-living individuals, was higher in all three regimes than in flies on a constant diet of CD or PDD (Fig. 4D). Panel E summarizes the median and maximum 10% of the lifespan data. Regime 1 also extended the median lifespan of control flies, with an increase of 47% compared to CD flies (Fig. 4F), and their maximum lifespan was extended by as much as 86% (Fig. 4E, 4F), which is considerably larger than the conventional lifelong mild protein restriction.

**Fig. 4:**
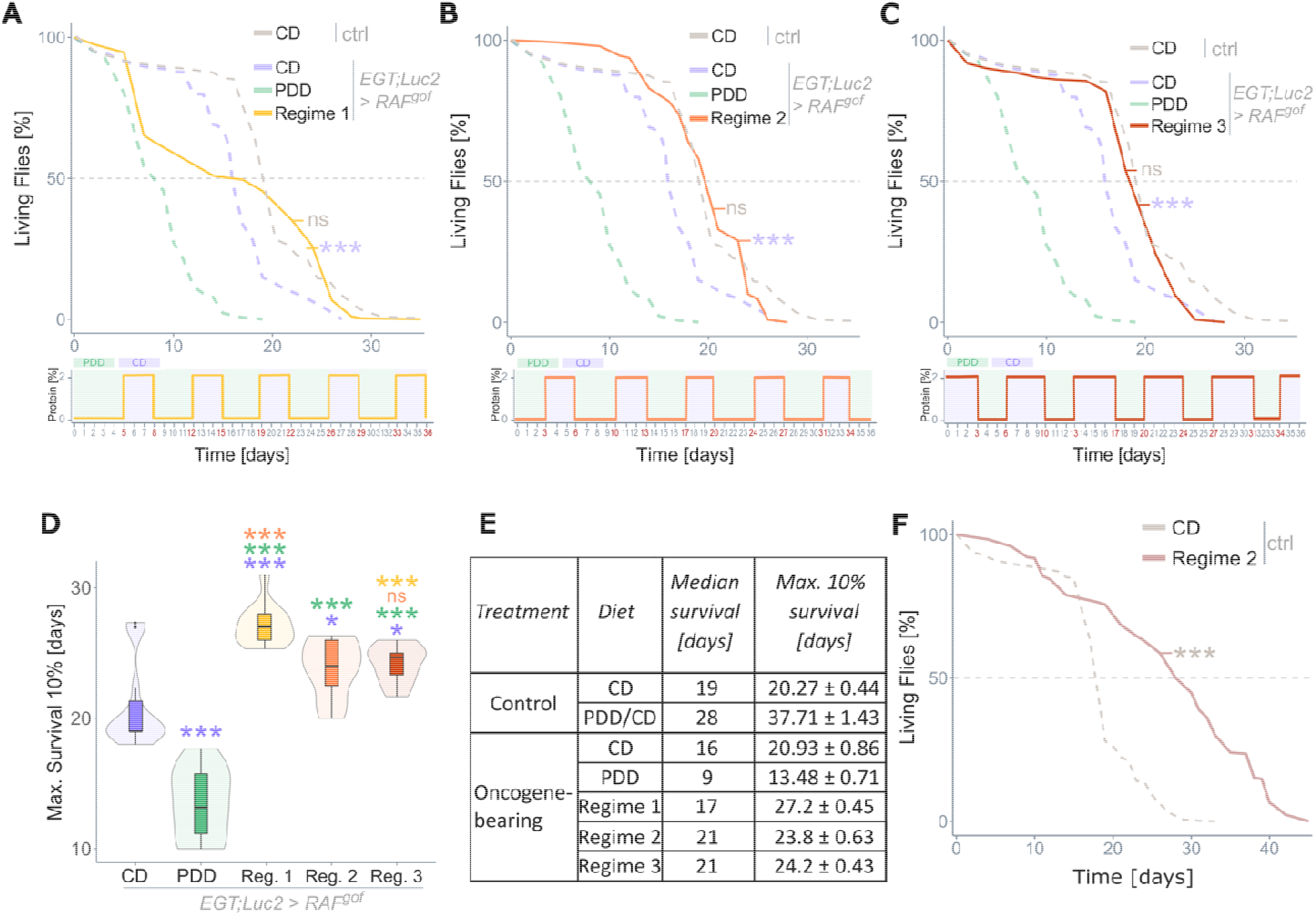
Recurrent severe protein depletion extends the lifespan of tumor-bearing flies. (A-C) Lifespans of RAF^gof^ expressing flies on various feeding regimes at 29°C, with details of each regime depicted below the plots. Regime 1 (A) and Regime 2 (B) only vary in the duration of the initial PDD phase. Regime 3 (C) starts with CD instead of PDD and had shorter alternating PDD and longer CD phases. All experiments were conducted simultaneously, and data are superimposed onto the data shown in Fig. 3A (dashed lines). (D) Maximum lifespans of the 10% longest-living flies. (E) Table summarizing median and maximum (10%) lifespans of all tested groups. (F) Lifespans of control flies subjected to Regime 1 or CD. All flies were kept at 29°C with n=360-450 flies per treatment.

### Recurrent protein depletion rescues tumor phenotype and reinstates gut physiology

After the initial phase and one full cycle of Regime 2, the tumor mass of the flies was assessed by microscopy of the thorax and abdomen and quantification of progenitor cells expressing luciferase. The cell layer of esg^+^ progenitor cells that formed in Raf^gof^ expressing flies fed on CD was much weaker after 10 days of Regime 2 feeding, with a phenotype like that of control flies. Cellular accumulation in both the thoracic and abdominal intestines was absent (Fig. 5A). The oncogene co-expressed luciferase activity increased during feeding on CD, but decreased during the PDD phase, as shown in Fig. 5B. Additional cross-sections of the anterior R2 midgut region are shown in Fig. S5. As all three tested recurrent feeding regimes rescued the lifespan of oncogene-expressing flies, further investigations of physiological and morphological outcomes were performed using Regime 2 only. The intestinal physiological parameters of the gut area per cross-section, lumen volume, and lumen/gut ratio, which were affected by tumor induction on CD, were all partially rescued by Regime 2 feeding (Fig. 5C-E). The fully nutritious CD diet increased the intestinal area in RAF^gof^-expressing flies and resulted in both absolute and relative obstruction of the intestine, as reflected by changes in the lumen volume and lumen-to-gut ratio, respectively. However, this phenotype was largely rescued by a recurrent protein-restricted feeding regime (Fig. 5D, E). The reinstated gut morphology was also reflected in the digestive function, as measured by fecal output. Even though measurements were taken after a 4-days period of PDD feeding (on day 10 of Regime 2), fecal spot size was rescued (Fig. 5F).

**Fig. 5:**
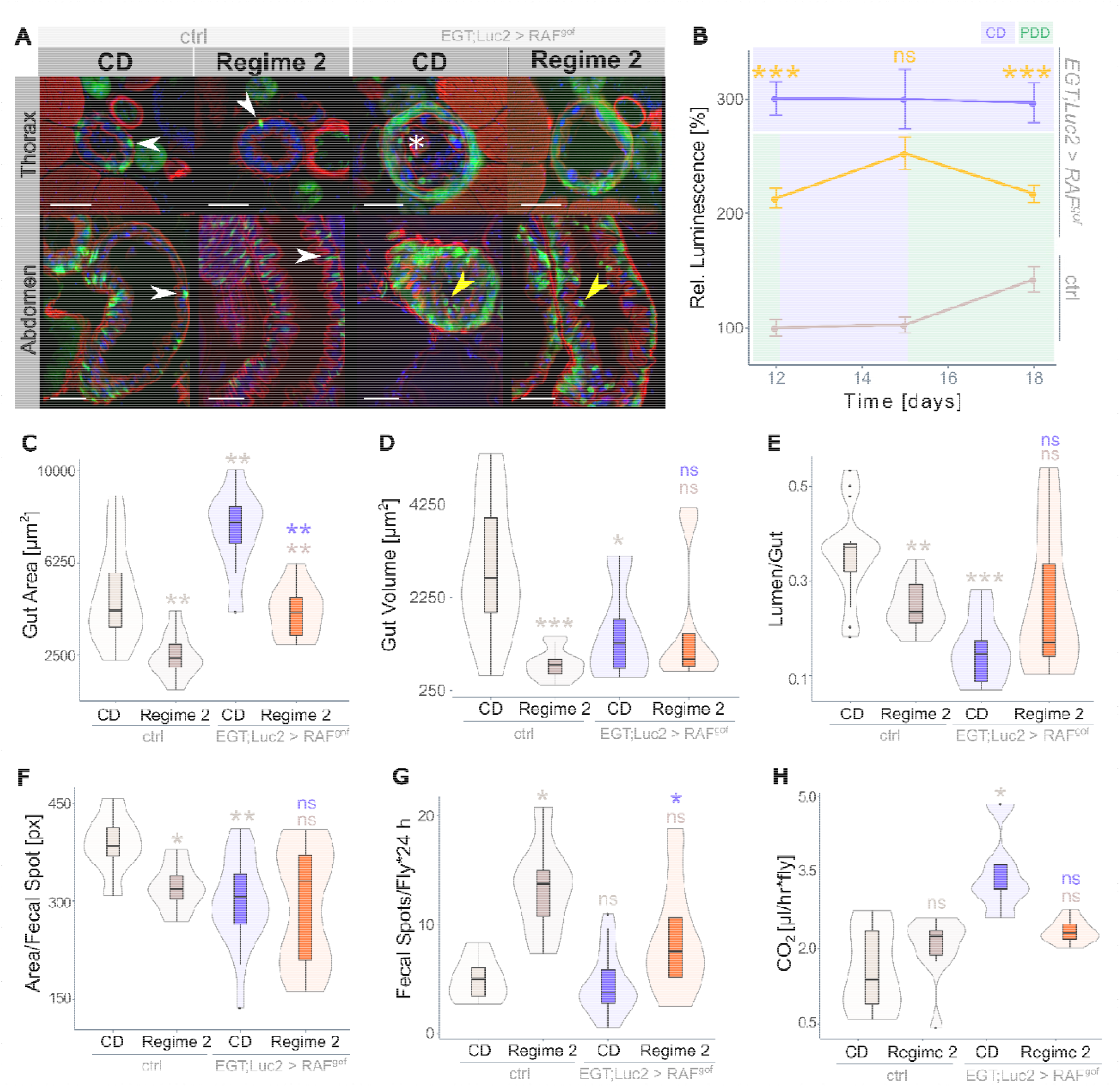
Recurrent protein depletion rescues tumor phenotype. (A) Sections through thorax (upper panel) or abdomen (lower panel) of control and RAF^gof^ expressing flies fed on CD and on Regime 2. The GFP^+^ esg^+^ cells attached to the basal membrane (arrowheads) and protruding mass (asterisk) visible in RAF^gof^ expressing flies subjected to CD are mostly absent following Regime 2 feeding. That feeding also resulted in detached esg^+^ cells (yellow arrowhead, scale bar: 50 μm). Micrographs were taken after 10 days of treatment. (B) Co-expressed luciferase in progenitor cells in control flies on CD and RAF^gof^ expressing flies on Regime 1 feeding for 12 to 18 days (n=5-10). (C) Total gut area per cross section, (D) lumen volume per cross section and (E) lumen-to-gut ratio of control and RAF^gof^ expressing flies subjected to CD or Regime 2 feeding for 10 days (n=9-11). (F) Fecal spot size and (G) fecal spot numbers at day 10 (n=7-21). (H) Basic metabolic rate at day 10 (n=5-10).

Together with a normal fecal spot abundance (Fig. 5G), this indicates a recovery of intestinal functionality in oncogene-bearing flies through a recurrent diet. Although these flies exhibited a two-fold increase in CO_2_ production after ten days of tumor induction when fed CD, this was restored to normal levels in Regime 2 (Fig. 5H). This lack of elevated resting metabolism was in concordance with the rescued morphological and functional intestinal phenotypes.

### Tumor growth overrides food-related shifts in intestinal microbiota composition

In humans, a shift in the diversity of the intestinal microbiota has been observed in patients with colorectal cancer, along with the establishment of individual taxa, such as *Fusobacterium nucleatum* which are not common in a healthy microbiome (51). When the intestinal microbiome of the flies was investigated by 16S DNA profiling, distinct clustering of the microbial communities of RAF^gof^ expressing and control flies was observed (Fig. 6A). The tumor phenotype induced by RAF^gof^ expression in ISCs and enteroblasts resulted in shifts in the intestinal microbial composition, resulting in a relative decrease in *Enterococceae* and *Streptococceae* and an increase in Beta- and Alphaproteobacteria compared to control flies (Fig. 6B). Interestingly, the microbiome was not affected by diet; neither control flies nor RAF^gof^ expressing flies subjected to CD, PDD, or Regime 2 resulted in further clustering of the obtained 16S sequences (Fig. 6C). In healthy organisms, stable, diet-dependent microbial shifts are usually established within a few days of a new feeding regime (52). However, this effect appeared to be overridden by the expression of the RAF^gof^ oncogene, as similar microbial changes were observed across all feeding regimes, with the microbial shift attributable to RAF^gof^ induction rather than dietary intake (Fig. 6D). The most noticeable and consistent changes were the enrichment of *Burkholderiales* and *Lactobacillaceae* together with a decrease in *Enterococceae* in oncogene-expressing flies compared with controls. Notably, the enrichment of *Burkholderiales* members is also frequently observed in patients with CRC (53), indicating a potentially highly conserved role for these bacterial species in intestinal tumor development.

**Fig. 6:**
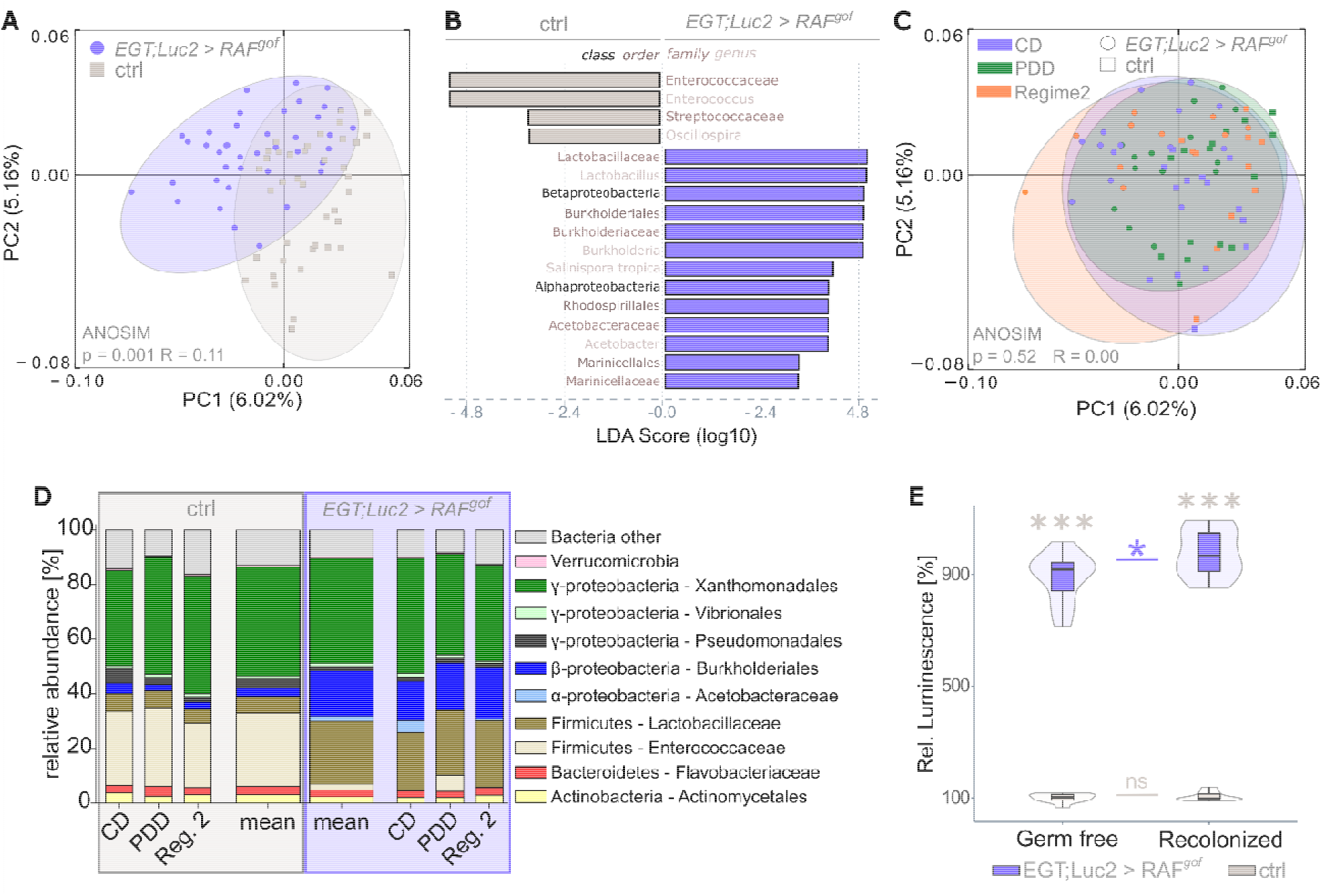
Tumor induction leads to a shift in microbiome composition that is independent of nutritional input. The intestinal microbiome was assessed by 16S rRNA analysis after 3, 6 and 10 days of tumor induction on the respective feeding regime (n=5 per each) and these timepoints were pooled prior to analysis. (A) Principal Coordinates Analysis (PCoA) segregates the sequence data from control and tumor-bearing flies. Ellipses were added manually. (B) Linear discriminant analysis (LDA) on relative abundance of taxonomic groups in the control and tumor-bearing flies. Bars represent taxa specifically associated with either genotype. (C) PCoA as in panel A, now colored for feeding regime. (D) Bar plots showing relative abundance of bacterial taxa obtained with the indicated diets and the mean of all data combined. (E) Luciferase activity in progenitor cells in control and oncogene-expressing flies raised germ free and following recolonization, after 3 days of oncogene induction on CD (n=10).

The intestinal microbiota plays a major role in organismal health, is responsible for food digestion and absorption, and, together with nutritional input and gut epithelial homeostasis, affects the host metabolism. Therefore, we tested the effect of the intestinal microbiota on the overall tumor load in axenically raised RAF^gof^ expressing flies and in such flies after recolonization with a microbial community mimicking a natural population. The abundance of intestinal progenitor cells in control flies remained unchanged because of recolonization; however, in oncogene-bearing flies, recolonization increased the tumor load compared to that in germ-free flies (Fig. 6E). This suggests that intestinal bacteria are not essential for tumor development but contribute to this process.

### Recurrent protein restriction mimics continuous protein limitation by transcriptomic changes that have a memory effect

To date, there is no satisfactory mechanistic explanation of how dietary regimes with alternating phases of calorie or protein restriction elicit beneficial effects. We hypothesized that protein restriction would create a memory effect that extends its positive effects beyond the actual PDD phase. This effect is most likely caused by transcriptional changes that persistently alter the gene expression. To test this hypothesis and to identify such persistent transcript signatures, transcriptome analysis was performed on midgut tissues from control and oncogene-expressing flies fed CD, PDD, and Regime 2. Principal Coordinate Analysis of the transcriptome data produced two clearly divided groups, separating oncogene-bearing flies from the controls (Fig. S6A) and further separating groups by feeding regime using principal component analysis (Fig. S6). Overall, the transcript levels of control flies were more similar between CD and Regime 2 than either were to PDD (Fig. S6B). A transcriptome heatmap (Fig. 7A) displayed genes with a resilient expression pattern (cluster 4) that returned to their original (CD) transcript levels after a PDD phase, resulting in similar expression levels between CD and Regime 2, but distinct from PDD (red box, Fig. 7A). In contrast, other gene clusters were upregulated or downregulated following PDD, but their expression remained at these levels during fully nutritious feeding in Regime 2, suggesting a memory effect. These are represented by clusters 2, 10, 11, and 19 in the heat map (green boxes, Fig. 7A). Focusing on these genes might reveal the key genes that convey the memory effect that enables recurrent protein-restricted feeding to elicit long-term PDD-like health benefits.

**Fig. 7:**
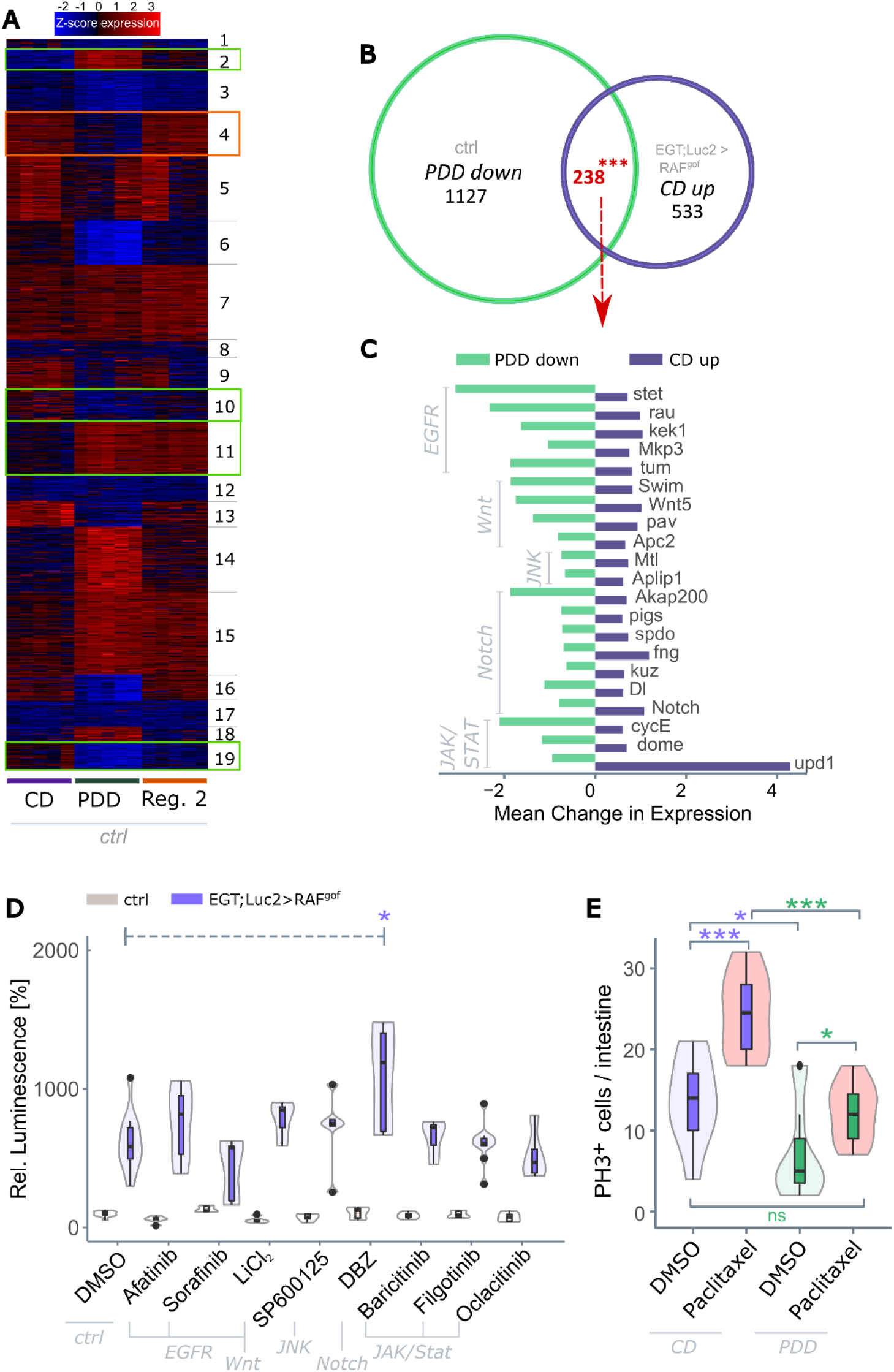
PDD has a distinct transcriptomic signature that can not be mimicked through drug exposure. (A) K-means clustering of differentially expressed genes (degs) of control flies feeding on control diet (CD), protein depleted diet (PDD), or the Regime 2 diet (cf. Fig. 4) for 13 days with p < 0.001 and fold change > 4 (n - 5). Clusters were identified that exhibited either a memory effect (green boxes, cluster 2, 10, 11, 19), or showed resilience (red box, cluster 4). (B) Venn diagram of degs that were downregulated in healthy flies on PDD and upregulated in tumor-bearing flies on CD (both compared to control flies on CD) after 13 days of oncogene induction (n=5). (C) Fold expression changes of a subset of the 238 degs identified in panel B that were involved in five major signaling pathways. (D) RAF^gof^ co-expressed luciferase signal in control flies and in flies expressing RAF^gof^ after 3 days of tumor induction and simultaneous subjection to pathway-specific inhibitors. Luminescence in healthy flies after 3 days at 29°C on DMSO is taken as 100% (n=4-19). (E) Number of actively proliferating cells as measured by quantifying phospho-histone H3^+^ (PH3^+^) cells in healthy *w^1118^* flies after 3 days of subjection to Paclitaxel or DMSO (n=9-15). DBZ = Dibenzazepine, DMSO = dimethyl sulfoxide.

We further classify these ‘memory’ genes by comparing all genes from control flies that were up- or downregulated in PDD *vs*. CD, to the genes that were regulated in a similar manner in Regime 2 *vs*. CD (Fig. S7). Of the 605 differentially expressed genes (degs) that were upregulated in the Regime 2 diet compared to the CD diet, 387 (64%) produced a similar pattern to that of the PDD diet. Likewise, of the 468 genes that were downregulated after Regime 2 compared to CD, 62% (290) were also downregulated with PDD. A total of 677 ‘memory’ genes were identified. GO term and KEGG pathway analyses of these genes revealed enriched terms in a variety of metabolic pathways (e.g., peptidase/lipase activity and thiamine metabolism) as well as responses to extrinsic stressors and immune stimulation. An excerpt of the enriched terms for genes with the strongest transcriptional shifts is shown in Fig. S7.

Next, we specifically concentrated on genes whose transcription was induced by tumor development (upregulated in tumor-bearing flies compared to control flies when both were fed CD) but simultaneously downregulated by protein restriction (downregulated on PDD compared to CD in control flies). These degs with antithetic expression patterns are particularly intriguing because they bear tumor development-associated transcript signatures that can be specifically reverted by recurrent dietary protein depletion. A total of 238 genes meeting these requirements were identified (Fig. 7B). GO analysis of these degs identified associations with cell division and proliferation events, which was expected, but also with genes involved in the cytoskeleton and cell structure (Table S1). The 238 degs of interest contained 21 genes whose products were involved in the EGFR, Wnt, JNK, Notch, and JAK/STAT signaling pathways. The amplitude of antithetic expression changes is shown in Figure 7C. The strongest downregulation with PDD feeding was observed for stet (EGFR pathway), *AKAP200* (Notch), and *cycE* (JAK/STAT), whereas the strongest upregulation in tumor development was observed for *upd1* (JAK/STAT pathway).

### PDD effects are not mimicked by drugs targeting developmental signaling pathways, but PDD prevents secondary stem cell proliferation

*We* investigated whether pharmaceutical intervention of developmental pathways could suppress tumor induction in response to RAF^gof^ overexpression. Oncogene-bearing flies were treated with commercially available pathway-specific inhibitors and their tumor loads were determined. However, none of the investigated inhibitors suppressed progenitor cell abundance on their own (Fig. 7D). Multiple combinations of inhibitors of various pathways were also tested; however, no effect was observed (Fig. S8). It is possible that various pathways act in a redundant manner, and that their simultaneous inhibition is required to produce an effect.

Chemotherapeutic treatment targets the elimination of actively proliferating tumor cells; however, it can induce unwanted secondary effects that induce stem cell activity and facilitate chemoresistance and metastasis (54, 55). In view of the tumor-suppressive activity of PDD, we investigated whether PDD could suppress such secondary outcomes. To this end, we monitored the abundance of actively proliferating stem cells after treatment with paclitaxel, a chemotherapeutic commonly used to manage multiple types of cancers that have been shown to induce secondary stem cell proliferation in *Drosophila* (31). The number of actively proliferating stem cells, marked by PH3 expression, was determined in the healthy control flies before and after exposure to paclitaxel (Fig. 7E). As expected, CD-fed flies had an increased abundance of PH3^+^ stem cells following paclitaxel exposure, whereas this effect was completely diminished in flies fed PDD during drug exposure. This indicates that subjection to PDD during tumor treatment does not induce excessive intestinal proliferation but rather resembles a catch-up effect that reestablishes normal levels of stem cell proliferation. This observation further indicates the therapeutic potential of an intermittent, severely protein-restrictive diet.

## Discussion

Growing tumors have special energy and metabolite requirements both quantitatively and qualitatively. Strategies that interfere with these specific needs are particularly promising for limiting tumor growth, particularly diets that restrict energy intake or supply specific macronutrients either continuously or intermittently (14, 56, 57). The individual effects of energy restriction, macronutrient restriction, and temporary or long-term reprogramming have not been completely disassembled (58). Dietary protein restriction synergistically limits tumor growth by reducing metabolite availability and reprogramming tumor cell metabolism (16, 59). This is consistent with the general health-promoting effects of protein restriction in humans and translates into life extensions in several model organisms (60, 61). Here, we used a *Drosophila* CRC model to establish that the inhibition of tumor growth correlates with the severity of dietary protein restriction. Severe protein restriction substantially reduces tumor burden in adult flies, reduces fat and muscle tissue, and shortens their lifespan. Severely reduced energy and metabolite intake lead to depletion of the body’s energy reserves and eventually death (62). Tissue wasting is a problem in many cancers and is involved in over 20% of all cancer-associated deaths (63). For therapeutic intervention, tumor growth reduction owing to nutritional restriction must be uncoupled from energy depletion to prevent negative outcomes. This was established by short phases of severe protein restriction, alternated with phases of a fully nutritious diet. It had already been shown that phases of moderate protein restriction alternated by a normal diet extends the lifespan of *Drosophila* similarly to continuous dietary restriction (27). This strategy was modified to apply the recurrent phases of severe protein reduction. The tested diet combined the anti-tumorigenic effects of strict protein depletion with increased longevity, and therefore has high potential as a sustainable anti-cancer diet. Three recurrent diets were tested, all containing phases of at least three days of protein restriction, as it was shown that this is needed for health-promoting effects (27). Although all three diets extended the median lifespan of the flies, an initial prolonged phase of protein depletion resulted in an initial higher death rate. This could be overcome by a shorter initial protein depletion period of three days, followed by four days of a fully nutritious diet, after which these phases were alternated. This feeding regime resulted in reduced tumor development to the same extent as prolonged protein restriction without fatal side effects. This recurrent diet normalized the fecal output of the flies, which was indicative of regular bowel movement and normal digestive activity. Digestion efficiency is a major indicator of intestinal malignancies, as tumor growth not only spatially restricts the intestine but can also result in malabsorption of nutrients (64).

It has been assumed that a dietary alteration of protein restriction and normal nutrition can at best extend lifespan to the degree of lifelong protein restriction (27). However, in our model, the recurrent diet was far superior to continuous protein restriction, extending the lifespan of oncogene-expressing flies compared with that of healthy control flies. Thus, severe protein restriction is feasible when interspersed with phases of normal feeding, an approach that has thus far only been tested for a strict reduction in caloric intake (65, 66). Strategies that alternate short periods of complete starvation (zero-calorie intake) with periods of a normal diet have been tested as adjunctive cancer therapies (67, 68). More recently, fasting phases have been enabled by means of a fasting-mimicking diet with very low calorie content (65, 69). These findings suggest that severe intermittent protein restriction may be as effective as or more effective than a fasting-mimicking diet.

The mechanisms underlying cyclic dietary restriction remain unclear. Such mechanisms must explain the long-lasting effects of short phases of restriction. Transcriptome analysis identified differentially expressed genes, whereby both tumor induction and feeding regimes led to independent and strong changes. Several genes were antipodally regulated, with upregulation of tumor induction and downregulation because of protein restriction. These genes point to a nodular junction that integrates oncogenic and nutrient-dependent metabolic regulation via major signaling pathways. These included genes involved in Notch, JAK/STAT, JNK, and EGFR pathways. Their potential as pharmacological interventions was investigated, but none of the tested pathway-specific inhibitors had an effect on tumor development, either alone or in combination. Clearly, the molecular effects of protein limitation are highly complex, in line with the extensive transcriptome changes reported here, the downstream effects of which most likely interact at the organismal level. Beneficial dietary restrictions are unlikely to be attributable to a single target (70). This also implies that selection of escape mutations that might overcome the effects of a recurrent protein-restrictive diet is unlikely.

The effects of tumor induction and diet on the intestinal microbiota revealed that oncogene expression, but not diet, affected the composition of the bacterial community, producing major shifts in the relative abundance of bacterial taxa. Dietary changes are known to result in microbial shifts in the intestine (52). However, this effect was overwritten by tumor induction, leading to consistent shifts in microbial abundance in Raf^gof^ expressing flies independent of diet or tumor mass development. This finding is consistent with recent results showing a vicious circle between tumor-induced epithelial dysfunction and the development of dysbiosis (71). Intestinal functionality is highly dependent on the microbial community for nutrient breakdown, and availability and shifts can result in deleterious dysbiosis (72–76). The intestinal microbial composition of healthy and oncogene-expressing flies revealed distinct clustering, with a shift towards dysbiotic microbiota. Certain bacteria have been shown to regulate intestinal homeostasis by promoting proliferation via JAK-STAT and JNK activation in healthy flies (77). Two key members of the *Drosophila* intestinal microbiota, *Lactobacillus plantarum* and *Acetobacter pomorum*, can promote tissue growth by activating insulin signaling and accessing the TOR signaling pathway (78). We confirmed that the presence of microbiota affects tumor development, as fewer tumorigenic progenitor cells were present in axenically raised flies than in flies reconstituted with controlled commensal microbiota. The bacterial mix used for reconstitution contained *L. plantarum* and *A. pomorum*, and the increased tumor load in recolonized flies may be attributable to their growth-promoting effects or to an increased inflammatory status, which in turn exacerbates tumor growth. Specific members of the intestinal microbiota, such as *F. nucleatum*, act as catalysts for tumor progression (79) however, this species was not enriched in oncogene-expressing flies.

An undesirable effect of chemotherapy is induction of secondary stem cell activation and metastasis. Chemotherapeutics halt the proliferation of stem cells needed for tissue renewal, as well as that of tumor cells, selecting for mutations that can lead to tumor re-establishment, drug resistance, and increased metastatic potential (31, 80, 81). We found that this secondary effect was eliminated by the recurrent severe protein-restricted diet, as demonstrated by paclitaxel treatment of healthy flies. This increased stem cell proliferation, but only on a normal diet, whereas no increase in proliferation was observed in the PDD subjected flies. This indicates a clear benefit of such a diet in the prevention of tumor relapse, although further research is needed to confirm this and provide mechanistic explanations. The substantial benefit of nutritional interventions to reduce tumor relapse was recently demonstrated with FMD in breast cancer treatment (82).

In summary, our study showed that ectopic expression of the RAF^gof^ oncogene in the ISCs and enteroblasts of *D. melanogaster* led to intestinal overproliferation and obstruction, which was rescued by a recurrent diet with alternating phases of strict protein depletion and refeeding on a fully nutritive diet. This diet reduced the tumor load, rehabilitated intestinal morphology and function, and prolonged the lifespan of the flies. Transcriptomic reprogramming has been used to identify genes involved in the reduction of tumor progression. Multiple signaling cascades are involved, which may explain why reprogramming cannot be mimicked by therapeutic intervention. The inhibition of tumor growth through the proposed recurrent feeding regime coexists with changes in multiple signaling pathways, rendering the emergence of escape routes and resistance mechanisms highly unlikely. The nutritional regime proposed here holds great potential for developing therapeutic strategies, not only targeting RAF-dependent intestinal cancers but also for a variety of other malignancies. Our findings represent a vital step toward a systemic approach to cancer therapy that can be pursued in combination with established cancer treatments.

## Competing interests

The authors declare no competing interests.

## Acknowledgments

We are thankful to Michelle Markstein for the generous gift of flies and Carlos Ribeiro for their kind gift of bacterial strains. This work was supported by the German Research Foundation (DFG, CRC 1182, project C2 and EXC 2167, RTF-VIII, INST 257/591-1 FUGG) and the BMBF (Project DroLuCa).

## Author contributions

RP designed the experiments and analyzed and interpreted the data. RP and TR wrote the first draft of this manuscript. BM analyzed microbial data. JB, HH, BM, CK, and TR contributed to the project planning and manuscript writing. DE performed RNA-seq data analyses. PR-supervised sequencing. RP, JvF, JB, and CF performed experiments. RP, BM, and JvF advised on the microbiome experiments and data interpretation. TR conceived and supervised the project and interpreted the data. All authors contributed to the manuscript completion.

## Data and material availability

All data needed to draw conclusions are presented in the paper or in the Supplementary Material. Sequencing data were deposited in the Sequence Read Archive (SRA) under project ID PRJNA482856, and RNA-seq data were deposited at NCBI GEO (GSE134485).

## Notes

### Competing Interest Statement

The authors have declared no competing interest.

## References

1. G. B. D. C. C. Collaborators, Global, regional, and national burden of colorectal cancer and its risk factors, 1990-2019: a systematic analysis for the Global Burden of Disease Study 2019. Lancet Gastroenterol Hepatol 7, 627–647 (2022).

2. J. Ferlay et al., Cancer statistics for the year 2020: An overview. Int J Cancer 10.1002/ijc.33588 (2021).

3. N. Keum, E. Giovannucci, Global burden of colorectal cancer: emerging trends, risk factors and prevention strategies. Nat Rev Gastroenterol Hepatol 16, 713–732 (2019).

4. N. Papadimitriou et al., Physical activity and risks of breast and colorectal cancer: a Mendelian randomisation analysis. Nat Commun 11, 597 (2020).

5. S. K. Veettil et al., Role of Diet in Colorectal Cancer Incidence: Umbrella Review of Meta-analyses of Prospective Observational Studies. JAMA Netw Open 4, e2037341 (2021).

6. J. Li, X. Ma, D. Chakravarti, S. Shalapour, R. A. DePinho, Genetic and biological hallmarks of colorectal cancer. Genes Dev 35, 787–820 (2021).

7. H. Raskov, J. H. Soby, J. Troelsen, R. D. Bojesen, I. Gogenur, Driver Gene Mutations and Epigenetics in Colorectal Cancer. Ann Surg 271, 75–85 (2020).

8. B. P. Leitner, S. Siebel, N. D. Akingbesote, X. Zhang, R. J. Perry, Insulin and cancer: a tangled web. Biochem J 479, 583–607 (2022).

9. B. G. Allen et al., Ketogenic diets as an adjuvant cancer therapy: History and potential mechanism. Redox Biol 2, 963–970 (2014).

10. V. D. Longo, R. M. Anderson, Nutrition, longevity and disease: From molecular mechanisms to interventions. Cell 185, 1455–1470 (2022).

11. O. Dmitrieva-Posocco et al., beta-Hydroxybutyrate suppresses colorectal cancer. Nature 605, 160–165 (2022).

12. P. Rous, The Influence of Diet on Transplanted and Spontaneous Mouse Tumors. J Exp Med 20, 433–451 (1914).

13. R. J. Colman et al., Caloric restriction delays disease onset and mortality in rhesus monkeys. Science 325, 201–204 (2009).

14. M. Alidadi et al., The effect of caloric restriction and fasting on cancer. Semin Cancer Biol 73, 30–44 (2021).

15. L. Fontana et al., Dietary protein restriction inhibits tumor growth in human xenograft models. Oncotarget 4, 2451–2461 (2013).

16. M. E. Levine et al., Low protein intake is associated with a major reduction in IGF-1, cancer, and overall mortality in the 65 and younger but not older population. Cell Metab 19, 407–417 (2014).

17. J. Yin, W. Ren, X. Huang, T. Li, Y. Yin, Protein restriction and cancer. Biochim Biophys Acta Rev Cancer 1869, 256–262 (2018).

18. S. Brandhorst et al., A Periodic Diet that Mimics Fasting Promotes Multi-System Regeneration, Enhanced Cognitive Performance, and Healthspan. Cell Metab 22, 86–99 (2015).

19. I. Y. Choi et al., A Diet Mimicking Fasting Promotes Regeneration and Reduces Autoimmunity and Multiple Sclerosis Symptoms. Cell Rep 15, 2136–2146 (2016).

20. C. W. Cheng et al., Fasting-Mimicking Diet Promotes Ngn3-Driven beta-Cell Regeneration to Reverse Diabetes. Cell 168, 775–788 e712 (2017).

21. V. D. Longo, M. Di Tano, M. P. Mattson, N. Guidi, Intermittent and periodic fasting, longevity and disease. Nat Aging 1, 47–59 (2021).

22. M. Wei et al., Fasting-mimicking diet and markers/risk factors for aging, diabetes, cancer, and cardiovascular disease. Sci Transl Med 9 (2017).

23. C. Vernieri et al., Fasting-Mimicking Diet Is Safe and Reshapes Metabolism and Antitumor Immunity in Patients with Cancer. Cancer Discov 12, 90–107 (2022).

24. J. Arends et al., ESPEN guidelines on nutrition in cancer patients. Clin Nutr 36, 11–48 (2017).

25. X. Gao et al., Dietary methionine influences therapy in mouse cancer models and alters human metabolism. Nature 572, 397–401 (2019).

26. O. D. K. Maddocks et al., Modulating the therapeutic response of tumours to dietary serine and glycine starvation. Nature 544, 372–376 (2017).

27. R. Romey-Glusing et al., Nutritional regimens with periodically recurring phases of dietary restriction extend lifespan in Drosophila. FASEB J 10.1096/fj.201700934R, fj201700934R (2018).

28. M. Sonoshita, R. L. Cagan, Modeling Human Cancers in Drosophila. Curr Top Dev Biol 121, 287–309 (2017).

29. D. Bilder, K. Ong, T. C. Hsi, K. Adiga, J. Kim, Tumour-host interactions through the lens of Drosophila. Nat Rev Cancer 21, 687–700 (2021).

30. E. Bangi et al., A Drosophila platform identifies a novel, personalized therapy for a patient with adenoid cystic carcinoma. iScience 24, 102212 (2021).

31. M. Markstein et al., Systematic screen of chemotherapeutics in Drosophila stem cell tumors. Proc Natl Acad Sci U S A 111, 4530–4535 (2014).

32. M. Rera et al., Modulation of longevity and tissue homeostasis by the Drosophila PGC-1 homolog. Cell Metab 14, 623–634 (2011).

33. A. S. Yatsenko, A. K. Marrone, M. M. Kucherenko, H. R. Shcherbata, Measurement of metabolic rate in Drosophila using respirometry. J Vis Exp 10.3791/51681, e51681 (2014).

34. M. T. Wayland et al., Spotting the differences: probing host/microbiota interactions with a dedicated software tool for the analysis of faecal outputs in Drosophila. J Insect Physiol 69, 126–135 (2014).

35. J. Ro, Z. M. Harvanek, S. D. Pletcher, FLIC: high-throughput, continuous analysis of feeding behaviors in Drosophila. PLoS One 9, e101107 (2014).

36. A. Proske, J. Bossen, J. von Frieling, T. Roeder, Low-protein diet applied as part of combination therapy or stand-alone normalizes lifespan and tumor proliferation in a model of intestinal cancer. Aging (Albany NY) 13, 24017–24036 (2021).

37. R. Leitao-Goncalves et al., Commensal bacteria and essential amino acids control food choice behavior and reproduction. PLoS Biol 15, e2000862 (2017).

38. P. Rausch et al., Comparative analysis of amplicon and metagenomic sequencing methods reveals key features in the evolution of animal metaorganisms. Microbiome 7, 133 (2019).

39. B. J. Haas et al., Chimeric 16S rRNA sequence formation and detection in Sanger and 454- pyrosequenced PCR amplicons. Genome Res 21, 494–504 (2011).

40. J. G. Caporaso et al., QIIME allows analysis of high-throughput community sequencing data. Nat Methods 7, 335–336 (2010).

41. N. Segata et al., Metagenomic biomarker discovery and explanation. Genome Biol 12, R60 (2011).

42. D. Kim et al., TopHat2: accurate alignment of transcriptomes in the presence of insertions, deletions and gene fusions. Genome Biol 14, R36 (2013).

43. S. Anders, P. T. Pyl, W. Huber, HTSeq--a Python framework to work with high-throughput sequencing data. Bioinformatics 31, 166–169 (2015).

44. M. I. Love, W. Huber, S. Anders, Moderated estimation of fold change and dispersion for RNA-seq data with DESeq2. Genome Biol 15, 550 (2014).

45. H. Wickham, ggplot2: Elegant Graphics for Data Analysis (Springer, New York, 2009).

46. R. C. Team, R: A language and environment for statistical ‡‡‡‡ computing (R Foundation for Statistical Computing, Vienna, Austria, 2021).

47. S. X. Ge, D. Jung, R. Yao, ShinyGO: a graphical gene-set enrichment tool for animals and plants. Bioinformatics 36, 2628–2629 (2020).

48. C. Wagner et al., Constitutive immune activity promotes JNK- and FoxO-dependent remodeling of Drosophila airways. Cell Rep 35, 108956 (2021).

49. M. Tatar, S. Post, K. Yu, Nutrient control of Drosophila longevity. Trends Endocrinol Metab 25, 509–517 (2014).

50. J. H. Catterson et al., Short-Term, Intermittent Fasting Induces Long-Lasting Gut Health and TOR-Independent Lifespan Extension. Curr Biol 28, 1714–1724 e1714 (2018).

51. Y. Cheng, Z. Ling, L. Li, The Intestinal Microbiota and Colorectal Cancer. Front Immunol 11, 615056 (2020).

52. L. A. David et al., Diet rapidly and reproducibly alters the human gut microbiome. Nature 505, 559–563 (2014).

53. Q. Feng et al., Gut microbiome development along the colorectal adenoma-carcinoma sequence. Nat Commun 6, 6528 (2015).

54. C. D’Alterio et al., New CXCR4 Antagonist Peptide R (Pep R) Improves Standard Therapy in Colorectal Cancer. Cancers (Basel) 12 (2020).

55. S. Acharyya et al., A CXCL1 paracrine network links cancer chemoresistance and metastasis. Cell 150, 165–178 (2012).

56. L. C. D. Pomatto-Watson et al., Daily caloric restriction limits tumor growth more effectively than caloric cycling regardless of dietary composition. Nat Commun 12, 6201 (2021).

57. S. Brandhorst, V. D. Longo, Fasting and Caloric Restriction in Cancer Prevention and Treatment. Recent Results Cancer Res 207, 241–266 (2016).

58. V. A. Acosta-Rodriguez, M. H. M. de Groot, F. Rijo-Ferreira, C. B. Green, J. S. Takahashi, Mice under Caloric Restriction Self-Impose a Temporal Restriction of Food Intake as Revealed by an Automated Feeder System. Cell Metab 26, 267–277 e262 (2017).

59. S. Brandhorst, V. D. Longo, Protein Quantity and Source, Fasting-Mimicking Diets, and Longevity. Adv Nutr 10, S340–S350 (2019).

60. G. A. Soultoukis, L. Partridge, Dietary Protein, Metabolism, and Aging. Annu Rev Biochem 85, 5-34 (2016).

61. C. M. Hill et al., FGF21 is required for protein restriction to extend lifespan and improve metabolic health in male mice. Nat Commun 13, 1897 (2022).

62. Y. Liu, P. Saavedra, N. Perrimon, Cancer cachexia: lessons from Drosophila. Dis Model Mech 15 (2022).

63. V. E. Baracos, L. Martin, M. Korc, D. C. Guttridge, K. C. H. Fearon, Cancer-associated cachexia. Nat Rev Dis Primers 4, 17105 (2018).

64. J. Keller, P. Layer, The Pathophysiology of Malabsorption. Viszeralmedizin 30, 150–154 (2014).

65. I. Caffa et al., Fasting-mimicking diet and hormone therapy induce breast cancer regression. Nature 583, 620–624 (2020).

66. L. Raffaghello et al., Starvation-dependent differential stress resistance protects normal but not cancer cells against high-dose chemotherapy. Proc Natl Acad Sci USA 105, 8215–8220 (2008).

67. S. de Groot, H. Pijl, J. J. M. van der Hoeven, J. R. Kroep, Effects of short-term fasting on cancer treatment. J Exp Clin Cancer Res 38, 209 (2019).

68. A. Nencioni, I. Caffa, S. Cortellino, V. D. Longo, Fasting and cancer: molecular mechanisms and clinical application. Nat Rev Cancer 18, 707–719 (2018).

69. S. de Groot et al., Fasting mimicking diet as an adjunct to neoadjuvant chemotherapy for breast cancer in the multicentre randomized phase 2 DIRECT trial. Nat Commun 11, 3083 (2020).

70. C. L. Green, D. W. Lamming, L. Fontana, Molecular mechanisms of dietary restriction promoting health and longevity. Nat Rev Mol Cell Biol 23, 56–73 (2022).

71. J. Zhou, M. Boutros, JNK-dependent intestinal barrier failure disrupts host-microbe homeostasis during tumorigenesis. Proc Natl Acad Sci U S A 117, 9401–9412 (2020).

72. E. Ragonnaud, A. Biragyn, Gut microbiota as the key controllers of “healthy” aging of elderly people. Immun Ageing 18, 2 (2021).

73. K. Takahashi et al., Reduced Abundance of Butyrate-Producing Bacteria Species in the Fecal Microbial Community in Crohn’s Disease. Digestion 93, 59–65 (2016).

74. X. Xu, X. Zhang, Effects of cyclophosphamide on immune system and gut microbiota in mice. Microbiol Res 171, 97–106 (2015).

75. F. Imhann et al., Proton pump inhibitors affect the gut microbiome. Gut 65, 740–748 (2016).

76. S. R. Modi, J. J. Collins, D. A. Relman, Antibiotics and the gut microbiota. J Clin Invest 124, 4212–4218 (2014).

77. N. Buchon, N. A. Broderick, S. Chakrabarti, B. Lemaitre, Invasive and indigenous microbiota impact intestinal stem cell activity through multiple pathways in Drosophila. Genes Dev 23, 2333–2344 (2009).

78. A. Arias-Rojas, I. Iatsenko, The Role of Microbiota in Drosophila melanogaster Aging. Front Aging 3, 909509 (2022).

79. C. Kong et al., Fusobacterium Nucleatum Promotes the Development of Colorectal Cancer by Activating a Cytochrome P450/Epoxyoctadecenoic Acid Axis via TLR4/Keap1/NRF2 Signaling. Cancer Res 81, 4485–4498 (2021).

80. G. S. Karagiannis, J. S. Condeelis, M. H. Oktay, Chemotherapy-Induced Metastasis: Molecular Mechanisms, Clinical Manifestations, Therapeutic Interventions. Cancer Res 79, 4567–4576 (2019).

81. C. D’Alterio, S. Scala, G. Sozzi, L. Roz, G. Bertolini, Paradoxical effects of chemotherapy on tumor relapse and metastasis promotion. Semin Cancer Biol 60, 351–361 (2020).

82. G. Salvadori et al., Fasting-mimicking diet blocks triple-negative breast cancer and cancer stem cell escape. Cell Metab 33, 2247–2259 e2246 (2021).

