## Supplemental Material for "Recurrent phases of strict protein limitation inhibit tumor growth and restore lifespan in a *Drosophila* intestinal cancer model"

### Recurrent phases of protein depletion rescue tumor phenotypes in a *Drosophila* intestinal cancer model

**This file includes Supplementary Figures S1 to S7 and Table S1**

**Fig. S1**

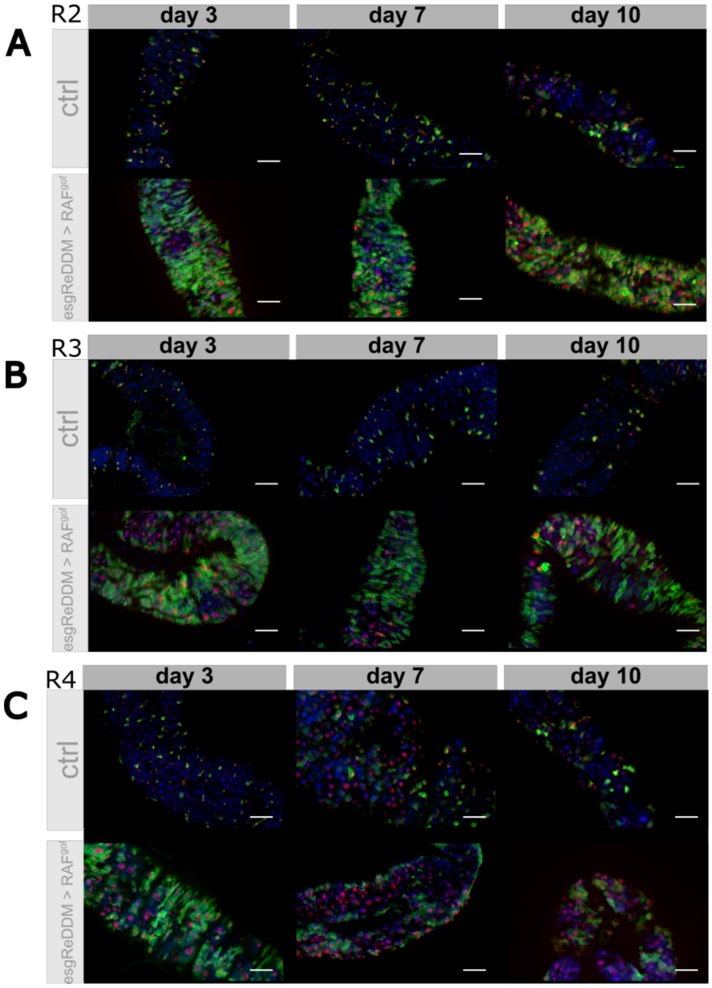


**Fig. S1: Lineage tracing in midgut region R2 (A), R3 (B), and R4 (C) of control flies and flies expressing RAF$^{gof}$ in ISCs and EBs.** The ReDDM system (1) was used to visualize the development of ISCs (expressing GFP) into enteroblasts (expressing RFP). Intestines were dissected after 3, 7 or 10 days of tumor induction from flies on control diet. During differentiation the short-lived GFP fades, leaving fully differentiated enteroblasts marked with RFP only (scale bar 50µm). ctrl = esgReDDM > w1118.

(1) Antonello ZA, Reiff T, Ballesta-Illan E, Dominguez M. Robust intestinal homeostasis relies on cellular plasticity in enteroblasts mediated by miR-8-Escargot switch. EMBO J. 201534(15):2025-2041.

**Fig. S2**

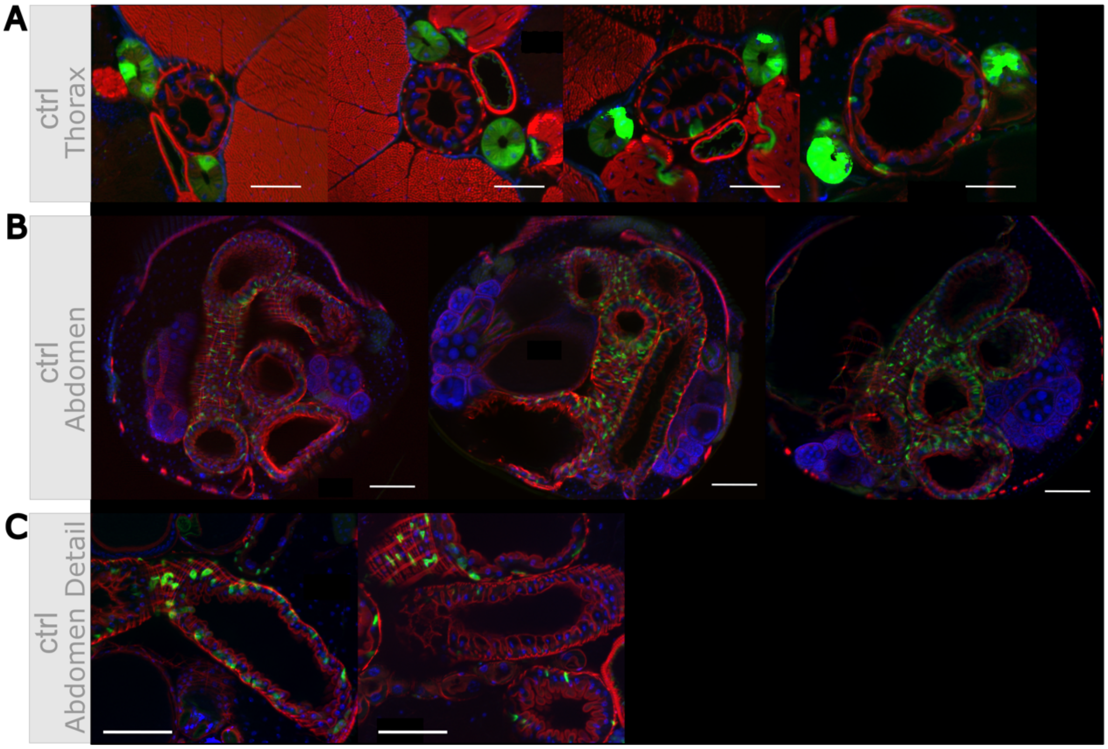


**Fig. S2: Intestinal structure and phenotype variability of healthy control flies and oncogene expressing flies on a nutritious control diet.** A-C, top: control flies (EGT;Luc2 > $w^{1118}$) and A-C, bottom: oncogene-bearing flies (EGT;Luc2 > $RAF^{gof}$ ) after 3 days of induction, showing (A) sections through the thorax (anterior midgut, R2) (scale bar 50 µm); (B) section showing multiple intestinal loops (R3-5 midgut region) (scale bar 100 µm); and (C) detail of an intestinal loop (region not specified, scale bar 100 µm).

**Fig. S3**

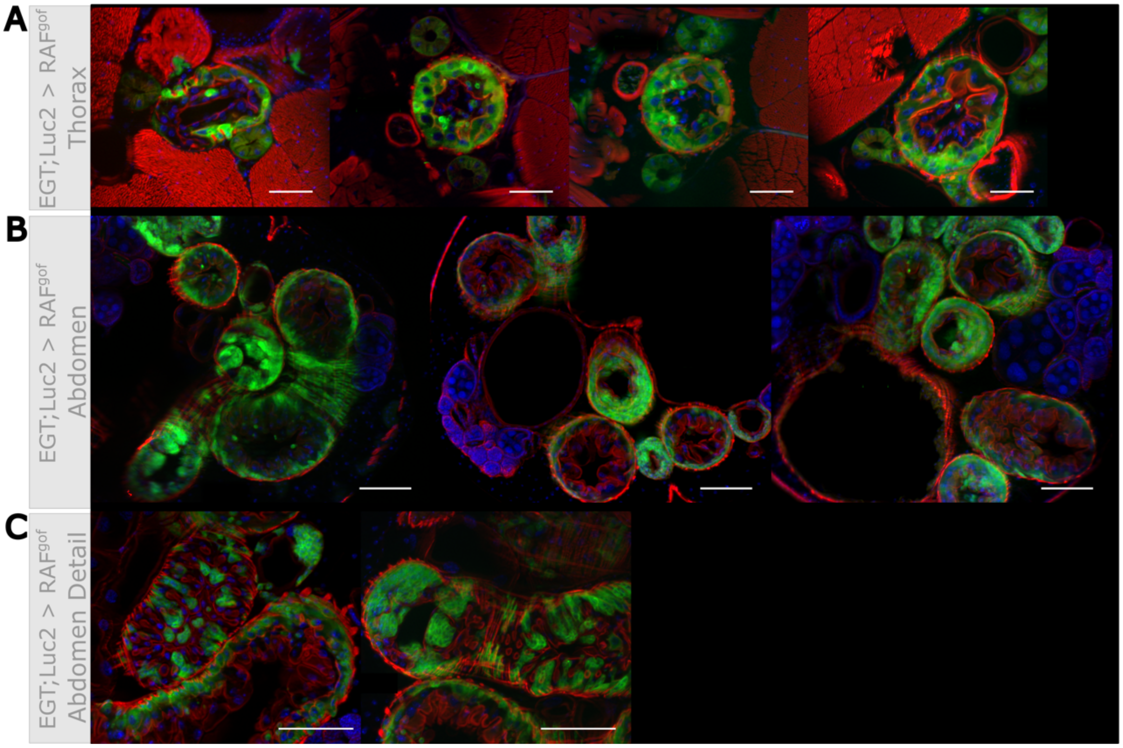


**Fig. S3: Intestinal structure and phenotype variability of oncogene-expressing EGT;Luc2 > RAF$^{gof}$ flies feeding on PDD.** (A) Sections through the thorax (anterior midgut, R2) (scale bar 50 µm). (B) Section through the abdomen (R3-5 midgut region) showing multiple intestinal loops (scale bar 100 µm). (C) Detail of intestinal loop (scale bar 100 µm). The flies were fed for three days on a protein-depleted diet containing 0.1% yeast extract.

**Fig. S4**

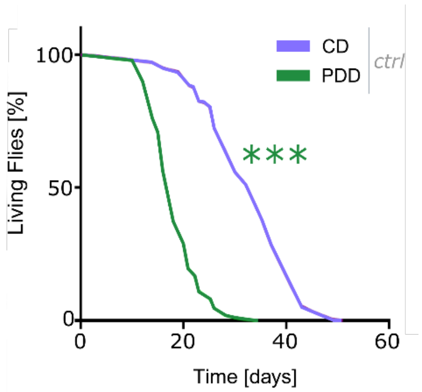


**Fig. S4:** Lifespan of control flies subjected to PDD or CD at 29°C (n = 200-205). ctrl = $w^{1118}$, CD = normal medium, PDD = protein depletion medium, *** $p < 0.001$.

**Fig. S5**

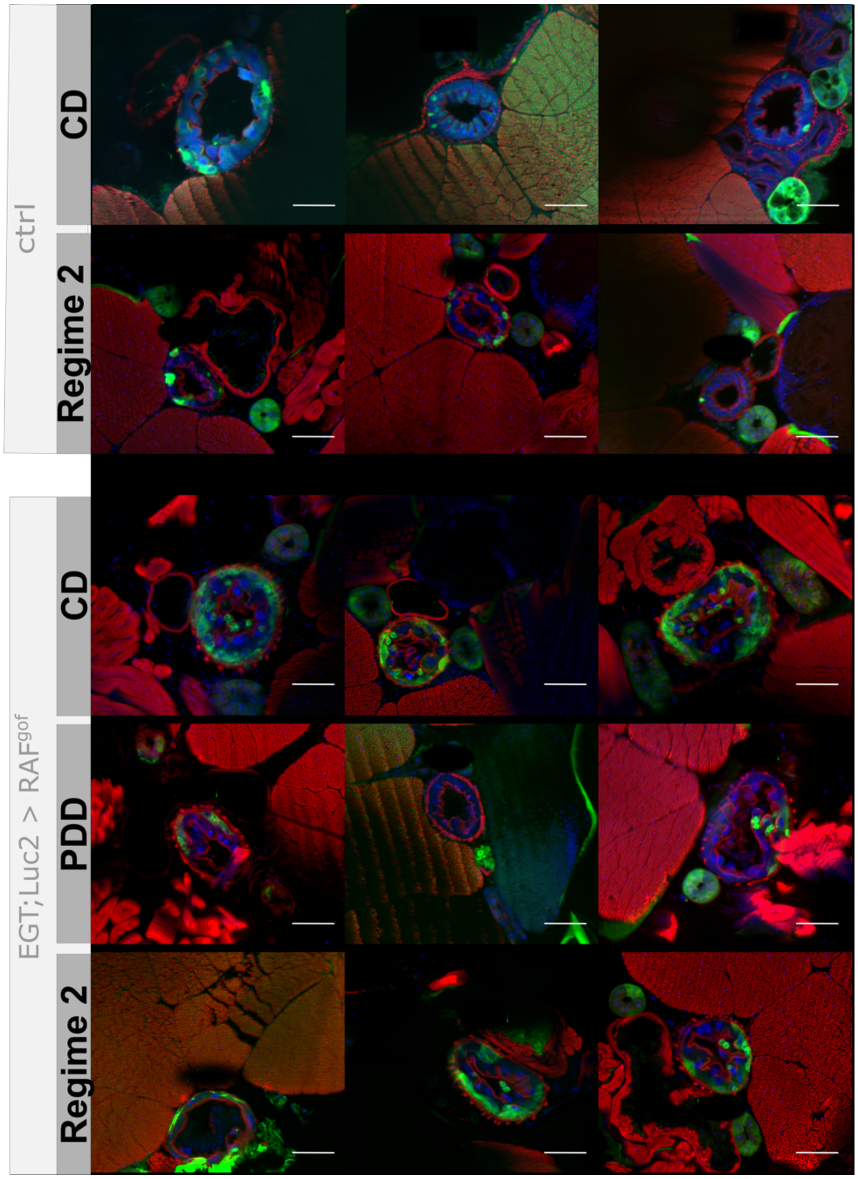


**Fig. S5:** Phenotypic observations in cross sections of the anterior R2 midgut region of flies feeding for 10 days on CD, feeding for 3 days on PDD followed by 3 days on CD and 4 days on PDD (PDD/CD) and 10 days on PDD (scale bar 50 µm). ctrl = EGT;Luc2 > $w^{1118}$, CD = control diet with nutritious medium, PDD = protein depleted diet.

**Fig. S6**

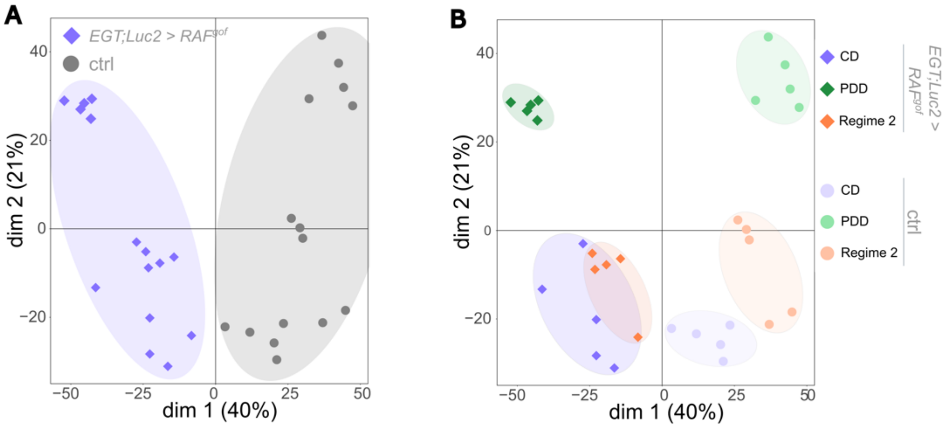


**Fig. S6: PCoA analysis of transcriptomic data.** Analysis of variation between and among samples using multi-dimensional scaling (MDS) was based on all transcripts (unweighted). The mRNA of oncogene-bearing and control flies was sequenced after 13 days of subjection to the respective feeding regime (n = 5). (A) Manually added ellipses represent tumor bearing (blue) and healthy controls (grey). (B) The same dataset as in A, now with ellipses representing feeding regimes as well as tumor or control phenotype (n = 5).

Fig. S7

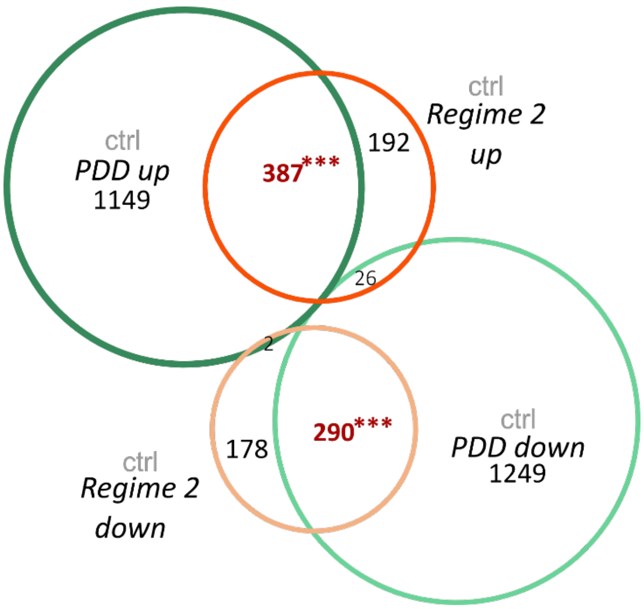


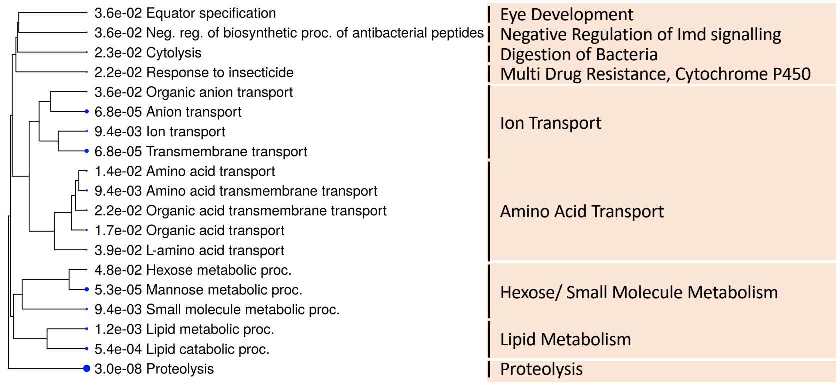


GO analysis of the cohort of genes upregulated in both Regime 2 and PDD

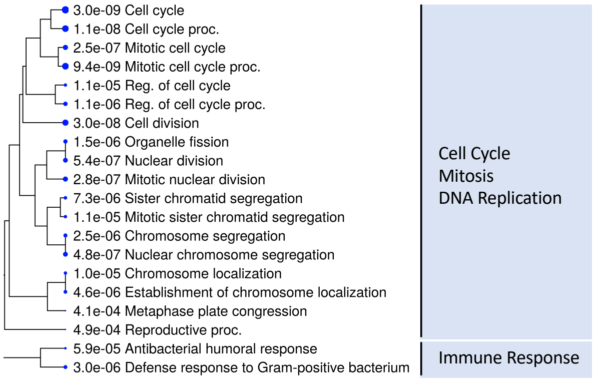


GO analysis of the cohort of genes downregulated in both Regime 2 and PDD

**Fig. S7**. Venn diagram identifying genes displaying a transcriptional memory effect (displayed in red). The GO terms of the commonly upregulated genes in Regime 2 and PDD (middle) and of the commonly downregulated genes (bottom) are shown as hierarchical clustering trees. The size of the solid circle corresponds to the enrichment FDR.

**Fig. S8**

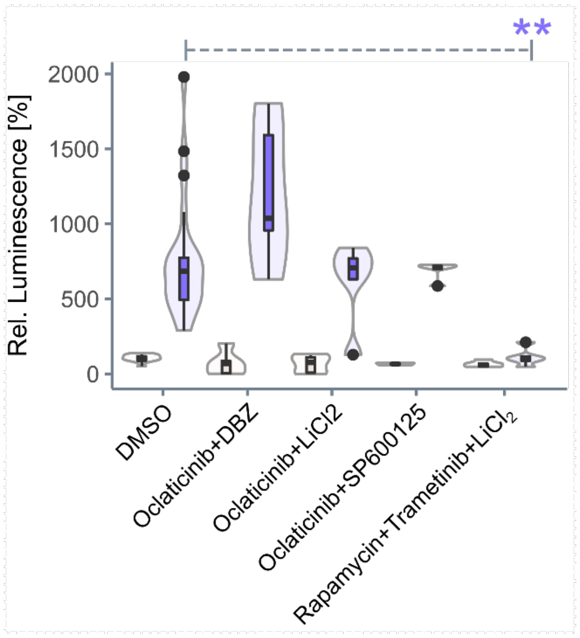


**Fig. S8: Multi-drug combinations do not mimic PDD.** RAF$^{gof}$ co-expressed luciferase signal was determined in control flies and in flies expressing RAF$^{gof}$ after 3 days of tumor induction. The flies were fed with CD containing the indicated combinations of pathway-specific inhibitors. is expressed with 100% for control flies given DMSO solvent. The combination of rapamycin, trametinib and $LiCl_2$ served as a positive control.

**Table S1:** Enrichment of GO terms for genes that were downregulated in healthy flies on PDD while upregulated in tumor-bearing flies on CD after 3 days of oncogene induction (n = 5).

| Term | Identifier | $log10^{-x}$ |
|---|---|---|
| cell division | GO:0051301 | 12.89 |
| mitotic cell cycle | GO:0000278 | 9.86 |
| spindle organization | GO:0007051 | 4.18 |
| DNA replication initiation | GO:0006270 | 3.70 |
| multicellular organismal reproductive process | GO:0048609 | 3.59 |
| cytoskeleton organization | GO:0007010 | 3.51 |
| cell differentiation | GO:0030154 | 1.94 |
| defense response to Gram-positive bacterium | GO:0050830 | 1.76 |
| anatomical structure development | GO:0048856 | 1.71 |
| embryonic organ development | GO:0048568 | 1.69 |
| actin filament -based process | GO:0030029 | 1.41 |
| animal organ development | GO:0048513 | 1.34 |
| DNA replication | KEGG:03030 | 3.42 |
| pentose and glucuronate interconversions | KEGG:00040 | 3.16 |
| ECM -receptor interaction | KEGG:04512 | 1.81 |
| retinol metabolism | KEGG:00830 | 1.63 |
| ascorbate and aldarate metabolism | KEGG:00053 | 1.49 |

Datasheet S1

Datasheet S2
